# Population-Level Disease Dynamics Reflect Individual Heterogeneities in Transmission

**DOI:** 10.1101/735480

**Authors:** Jonathon A. Siva-Jothy, Lauren A. White, Meggan E. Craft, Pedro F. Vale

**Affiliations:** Institute of Evolutionary Biology, School of Biological Sciences, University of Edinburgh, Ashworth Labs, Charlotte Auerbach Road, EH9 3JT Edinburgh, UK; SESYNC, 1 Park Place, Suite 300, Annapolis, MD 21401, USA; Department of Veterinary Population Medicine, University of Minnesota, St Paul, MN 55126

**Keywords:** *Drosophila melanogaster*, disease transmission, social aggregation, virus shedding, contact networks, disease modelling, transmission heterogeneity

## Abstract

Host heterogeneity in disease transmission is widespread and presents a major hurdle to predicting and minimizing pathogen spread. Using the *Drosophila melanogaster* model system infected with *Drosophila* C virus, we integrate experimental measurements of individual host heterogeneity in social aggregation, virus shedding, and disease-induced mortality into an epidemiological framework that simulates outbreaks of infectious disease. We use these simulations to calculate individual variation in disease transmission and apportion this variation to specific components of transmission: social network degree distribution, infectiousness, and infection duration. The experimentally-observed variation produces substantial differences in individual transmission potential, providing evidence for genetic and sex-specific effects on disease dynamics at a population level. Manipulating variation in social network connectivity, infectiousness, and infection duration in simulated populations reveals that these components affect disease transmission in clear and distinct ways. We consider the implications of this genetic and sex-specific variation in disease transmission and discuss implications for appropriate control methods given the relative contributions made by social aggregation, virus shedding, and infection duration to transmission in other host-pathogen systems.

## Introduction

Individual heterogeneity in host traits affecting disease transmission has major consequences for the predictability and severity of outbreaks of infectious disease, and in extreme cases can lead to ‘superspreaders’ or ‘supershedders’ of infection [1–4]. An individual’s transmission potential can be described as a function of its contact with susceptible individuals, the likelihood of that contact resulting in infection, and the length of time that individual remains infectious [5,6]. While the underlying causes of heterogeneity in transmission are poorly understood, each of these components may be affected by genetic variation in pathogen traits, the behavioural and physiological traits of the host, and their interaction with environmental factors [7–10]. While the effects of host contact behaviour on heterogeneity in pathogen transmission have been widely investigated [11–13], the role of variation in host physiological traits in generating heterogenous pathogen transmission are less clearly understood [6,14]. Moreover, the relative roles of these traits, and how they interact with one another in natural systems remain difficult to isolate and quantify.

One commonly used descriptor of how likely a disease is to spread through a population is the basic reproductive number, *R*_*0*_, which denotes the mean number of secondary cases caused by an infected individual in a susceptible population [5,15]. *R*_*0*_ is one the most widely used metrics in epidemiology, commonly used to predict outbreaks of infectious disease and as a theoretical tool to model pathogen evolution [16]. However, a potential shortcoming of *R*_0_ is that it reflects population averages, making it a poor predictor of disease outbreaks that arise due to extreme individual heterogeneity in transmission [17–19]. One way to address this shortcoming is to move beyond the population average transmission reflected in *R*_0_, and measure the *R*_0_ equivalent for each individual in that population, termed *V* [6,18,19]. This approach has the advantage of explicitly measuring the distribution of individual transmission (where the mean of the distribution is *R*_0_) and can be useful in identifying individuals at the most extreme of this distribution which could be likely superspreaders of infection. Another advantage of applying the basic reproduction number to individuals is that it provides an experimentally tractable framework to partition the variance in individual transmission among a range of behavioural, physiological and immune phenotypes that may lead to variation in *V* [6]

Notwithstanding these advantages, quantifying the behavioural, physiological and immune traits underlying the number of infections produced by a single individual remains tremendously challenging, particularly in wild or natural disease settings. One potentially useful approach is to measure *V* and its components by experimentally infecting model systems under controlled laboratory settings in order to quantify the roles of physiological and behavioural host heterogeneity on pathogen transmission [8,10,20,21]. This experimental approach offers the advantage of minimising environmental variation and allowing highly replicated measurements of individual host traits. However, such studies may be limited in their ability to extrapolate the effects of measured at the level of individual hosts to population-level epidemic dynamics. Mathematical modelling/*in silico* experiments are a useful tool to efficiently test different hypotheses in larger/scaled up populations and infer patterns across scales [22], but many theoretical studies often rely on assumptions about the level heterogeneity in host traits, in the absence of adequate empirical information [6,23]. The ideal approach is therefore to use mathematical modelling of epidemiological dynamics where as many parameters as possible are informed by experimental data measured on individual hosts in controlled laboratory settings.

Here we use this approach combining experiment and theory to test how population-level disease transmission dynamics are affected by empirically measured levels of variation in pathogen shedding, lifespan following infection and social aggregation. We previously measured individual-level variation in behavioural and physiological traits that are relevant to pathogen transmission in the fruit fly (*Drosophila melanogaster)* when infected with its viral pathogen *Drosophila* C Virus (DCV) [24,25]. These experiments leveraged genetic and sex-specific sources of variation in three traits that likely affect DCV transmission which occurs via larval or adult feeding [26]: the degree of group-level social aggregation (as an indicator of potential contact rate); mortality rate (which defines the duration of infection); and how much DCV each individual sheds into their environment (as a proxy measure of infectiousness).

We address three questions about the interactions of different types of behavioural and physiological heterogeneity in pathogen transmission. First, we asked if genetic and sex-specific variation in social aggregation, virus shedding, and duration of infection – as measured in lab setting – would result in different predicted epidemics in theoretical populations. In this initial set of simulations, theoretical populations were comprised of individuals with traits that were representative of the phenotypic heterogeneity in males or females of a single genetic background. By simulating and comparing epidemics in host populations comprised of a single sex and one genetic background, we focused on genetic and sex-specific sources of variation in disease transmission.

Second, to test the relative importance of index case genetic background and sex vs. group composition and how variation in transmission potential is affected by the diversity of the susceptible population, we simulated epidemics in populations where individuals’ traits are sampled from a larger phenotypic distribution, including males and females from ten genetic backgrounds. In these simulations, we varied the genetic background and sex of the index case.

Third, to test the relative importance of variation in specific host traits on epidemic dynamics, we compared epidemic dynamics of populations exhibiting empirically-measured levels of variation in social aggregation, viral shedding and mortality, to populations where we constrained variation in these traits to the population mean.

## Methods

### Measuring social aggregation, viral shedding and infection duration in infected *D. melanogaster*

Simulations were parameterised using experimental data on host aggregation, mortality, and viral shedding [24,25]. Readers are directed to that publication for a detailed description of data collection. Briefly, we established systemic infections with DCV in ten lines (Table 1) from the *Drosophila* Genetic Resource Panel (DGRP) (Mackay et al., 2012), chosen because they are extremes of susceptibility to DCV systemic infection [28]. DCV was cultured and prepared as described previously [29]. Systemic infections were achieved by pricking flies in the thorax near the pleural suture with a 0.5mm entomological needle dipped in DCV [30]. To measure lifespan following DCV infection, single flies were monitored daily until dead. DCV shedding was measured 1, 2, and 3 days after infection in 1.5ml Eppendorf tubes. Flies were removed from tubes after 24 hours and processed for qRT-PCR. DCV shed into the tube was collected by adding 100μl of TRI-reagent and thoroughly vortexing. TRI-reagent was removed for RNA extraction and reverse transcription before being quantified by qPCR [29]. To measure social aggregation, photos were taken of groups of 10-12 flies of the same genetic background, sex and infection status, in 55mm petri dishes. The mean number of neighbours each individual in a Petri dish had within circles of a 10mm, 15mm and 20mm threshold radius was calculated using the coordinates of each fly generated with the ImageJ multipoint tool.

**Table 1.** Factorial design and specifications for simulations testing the effect of genetic and sex-specific variation in social aggregation, viral shedding and susceptibility on population-level disease dynamics. We conducted 500 replicates per parameter set with 1000 individuals in the network. Simulations were allowed to run for 1000-time steps.

| Parameter | Levels |
| --- | --- |
| Genetic background | RAL-59, RAL-75, RAL-138, RAL-373, RAL-379, RAL-380, RAL-502, RAL-738, RAL-765, RAL-818 |
| Sex | Female, Male |
| Threshold radius ($r$) | 10mm, 15mm, 20mm |
| Pathogen transmission efficiency ($\tau$) | 0.1, 0.5, 1 |
| Scaled infectiousness ($\eta$) | 1, 2 |

### Simulation Methods

We used these empirical measurements from *D. melanogaster* to develop an individual-based, stochastic, static network model that tests how the sex-specific and genetic variation in viral shedding, susceptibility and social aggregation translates to differences in disease dynamics. Using a susceptible-infected (*SI*) model we simulated DCV transmission in a closed population with no births and where infected individuals can die [31]. Alongside empirically measured traits, we also tested the effect of pathogen viability and the relative infectiousness of supershedders. The effects of all parameters on outbreak dynamics were tested in a fully-factorial design. For each parameter set, 500 simulations were conducted for a population of 1000 individuals over the course of 1000 time steps (Tables 1, 2 & 3). Key metrics to measure outbreak dynamics included: fadeout likelihood, maximum number of infected individuals, outbreak duration, and time to maximum number of infected individuals. Code to conduct these simulations was written in *R* (Version 3.4.4) and is available at: https://github.com/whit1951/Drosophila

**Table 2.** Factorial design and specifications for simulations testing the effect of susceptible host diversity on disease transmission potential. We conducted 500 replicates per parameter set on networks of 1000 individuals (20 subgroups of 50 individuals each representing each sex/line combo). Simulations ran for 1000-time steps.

| Parameter | Levels |
| --- | --- |
| Index genetic background | RAL-59, RAL-75, RAL-138, RAL-373, RAL-379, RAL-380, RAL-502, RAL-738, RAL-765, RAL-818 |
| Index sex | Female, Male |
| Threshold radius | 10mm, 15mm, 20mm |
| Pathogen transmission efficiency ($\tau$) | 0.1, 0.5, 1 |
| Scaled infectiousness ($\eta$) | 1, 2 |

#### Social network degree distribution

To generate a simulated contact network reflecting contact rates of different phenotypes, we used a proxy for social aggregation: the number of neighbours within a set threshold radius. Individuals (nodes) within the prescribed threshold radius share an edge where transmission is possible. Using the number of neighbours within this radius for each fly, we derived a functional degree distribution for our simulated populations of interest. From this empirical degree distribution, we sampled 1000 times based on the number of individuals needed for the simulated network. This produced a network where the mean degree (rather than network density) was maintained between empirical and simulated populations. The dynamics of faecal-oral DCV transmission are poorly understood [32,33], but the virus is seen to readily proliferate through laboratory stocks of *Drosophila* [34]. To account for this and assess the relative importance of possible direct transmission routes, we consider the number of neighbours within 10, 15 or 20mm of one another and derive simulated social networks from these distinct degree distributions. Transmission is only possible between infected (*I*) and susceptible (*S*) flies within this set infectious distance. Importantly, using social aggregation as a proximate measure of contact rate assumes the likelihood of contact with DCV is proportional to an individual’s proximity to an infected fly.

#### Infectiousness

We estimated infectiousness (*κ*_*j*_) for any given infected individual, *j*, from our empirical measurements of viral shedding. The untransformed distribution of this data is highly skewed and zero-inflated, with some rare flies shedding exceedingly high viral titres (i.e., so-called supershedders), and others not shedding any virus at all (within the technical limit of detection). To account for this disparity, we used the natural log to transform our viral load shed distribution and divided these values by the greatest amount of virus shed, constraining infectiousness values between 0 and 1.

The amount of virus needed to ensure transmission is unclear. To account for this, we considered a ‘scaled infectiousness’ (*η*) parameter which had two levels, 1 or 2. This parameter reflects two hypotheses: (1) only supershedders at the upper end of our shedding distribution ensure 100% transmission, with all other individuals having a probability less than one; or (2) average and non-zero shedders could still shed enough to ensure infection, but supershedders increase the likelihood of transmission relative to average counterparts. The two levels of scaled infectiousness, 1 and 2, were implemented by multiplying our measure of infectiousness (*κ*_*j*_) by 1 or 2 respectively.

Another factor that may affect transmission is the viability of DCV in the environment without a host. To account for this, we included a transmission efficiency (*τ*) parameter into our model. The three levels, *τ* = 0.1, 0.5, or 1, altered infectiousness by multiplying the infectiousness value by 0.1, 0.5, and 1 respectively. The levels of transmission efficiency correspond to 10, 50, and 100% probability of transmission. Both scaled infectiousness (*η*) and transmission efficiency (*τ*) were held constant in simulations unless specifically mentioned.

#### Infection duration

DCV results in death for infected flies, making our empirical measurement of the time between inoculation and death an ideal measure of infection duration (*µ*). We calculated mortality rate as the inverse of empirical disease-related mortality for a given population. Once infected, individuals experienced a weighted coin flip probability of dying [B(1, 1/ *µ*)] at each time step.

#### Transmission rate

Combining all the elements above, transmission rate *β*_*ij*_ between an susceptible individual (*i*) and infectious host (*j*) is subject to the infectiousness of infectious host (*κ*_*j*_), the scaled infectiousness (*η* = 1 or 2), the transmission efficiency of the pathogen (*τ* = 0.1, 0.5 or 1), and whether or not an edge exists in the network between individuals *i* and 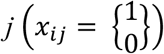:

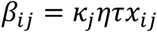

With individual-specific disease related mortality (*µ*_*j*_), transmission rate translates to differences in the number of susceptible (*S*) and infected (*I*) individuals at each time step according to:

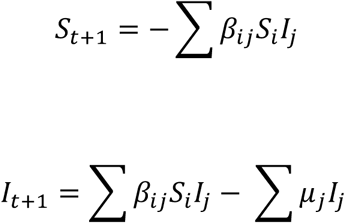

For each individual at each time step, infection and mortality were stochastic processes governed by Bernoulli draws, e.g., *Bernoulli*(*β*_*ij*_) and *Bernoulli*(*µ*_*j*_).

#### Theoretical simulation #1:The effect of genetic and sex-specific variation in social aggregation, viral shedding and susceptibility on pathogen transmission potential

We scaled-up empirical degree distributions for males and females of our ten genetic backgrounds to a theoretical population size of 1000. In each simulated population, flies were of the same sex and genetic background. We allowed infectiousness, duration of infection, and social aggregation to vary based on empirical measurements for each combination of sex and genetic background. For each individual simulation, we generated a new network from the scaled-up degree distribution, and randomly selected an individual from the network to start as the index case.

#### Theoretical simulation #2: The effect of population diversity on pathogen transmission potential

Many natural host populations have highly variable levels of diversity which can significantly affect host-pathogen dynamics [35]. To test if differences in transmission potential are robust to the diversity of the index case, we simulated populations where males and females of all ten genetic backgrounds were combined in equal proportion. More specifically, the simulated, scaled-up populations of 1000 individuals were comprised of 20 sub-populations each containing 50 sampled individuals drawn from the larger experimental distribution for each respective line/sex combo. Individuals maintained their respective empirical distributions for aggregation, infectiousness, and duration of infection according to their genetic background and sex combination. These simulated populations therefore reflect a relatively diverse population. We then varied which genetic background and sex combination served as the index case with 500 replicates per index case phenotype (Table 2).

#### Theoretical simulation #3: The consequences of variation in social aggregation, viral shedding and disease-related mortality on disease dynamics

To determine the relative importance of empirical variation in social aggregation, viral shedding, and disease-related mortality on disease transmission in a heterogeneous population, we simulated populations derived from the variation seen across all genetic backgrounds and both sexes. To determine the effect of population-level variation, we systematically constrained the variation in all three host traits to the population’s mean, individually and alongside one another. During these simulations, the unconstrained traits were free to vary according to the empirical measurements (Table 3). In the case of degree of the network, we rounded this value to ensure a whole number, which is essential for contact network formation (e.g., an individual cannot have 2.5 contacts). For example, to look at the effect of social aggregation by itself, we allowed social aggregation to take on the degree distribution of the entire heterogenous population but constrained viral shedding and infection duration to the empirically-measured means across all genetic backgrounds and both sexes.

**Table 3.** Experimental design for Experiment 3. We conducted 500 replicates per parameter set. Simulations ran for 1000-time steps.

| Parameter | Levels |
| --- | --- |
| Threshold radius | 10mm, 15mm, 20mm |
| Pathogen transmission efficiency ($\tau$) | 0.1, 0.5, 1 |
| Scaled infectiousness ($\eta$) | 1, 2 |
| Number of nodes in simulated network | 1000 |
| Vary social aggregation | TRUE, FALSE |
| Vary infectiousness | TRUE, FALSE |
| Vary infection duration | TRUE, FALSE |

### Random Forest Analysis

Parsing out the effects of individual variables in simulation modelling can be challenging because of collinear effects and sensitivity of frequentist measures of significance to sample size (White et al 2014). Random forest analysis is a machine learning approach that readily handles non-linear relationships between variables (Cutler et al., 2007). Here we utilize the *cforest* function from the *party* package in *R* to look at variable importance scores, which reflect the relative influence of each variable in the prediction of the random forest model [37,38]. The *party* package, in particular, addresses some of the potential biases of the original *randomForest* package that may result from continuous variables or variables with more categories [38]. For each theoretical experiment, we generated 1000 trees – at this level, no changes in variable importance order resulted from changes in the random seed suggesting a robust ranking order [38]. We reported variable importance scores as mean decrease in accuracy (a measure of permutation importance rather than node impurity), which describes the loss in accuracy resulting from randomly permuting the given variable [38,39].

### Outbreak Descriptors

We used five metrics to measure and characterise simulated outbreaks of infectious disease: fadeout likelihood, basic reproductive number (*R*_*0*_), maximum number of infected individuals, the time taken to reach the and maximum number of infected individuals, and outbreak duration. Fadeout likelihood represents the probability of an outbreak not occurring following the infection of the index case. It is the proportion of index cases that fail to transmit infection to at least one susceptible individual before dying from infection. We use *R*_*0*_ as a measure of the number of secondary cases of infection caused by the index case for the duration of the simulation.

## Results

As described in the Methods, in theoretical experiments 1 and 2, we tested full-factorial combinations of genetic background, sex, threshold radius, transmission efficiency and scaled infectiousness. In experiment 3, we tested full-factorial combinations of variation in infectiousness, social aggregation and infection duration, threshold radius, transmission efficiency and scaled infectiousness. Here, we present results with a threshold radius of 15mm, a transmission efficiency of 1, and a scaled infectiousness of 2 (Figure 1). We focussed on this combination of parameters to promote pathogen transmission in our simulations, while also allowing us to subsequently test the importance of direct transmission by comparing the 15mm threshold radius to 10mm and 20mm.

**Figure 1.**
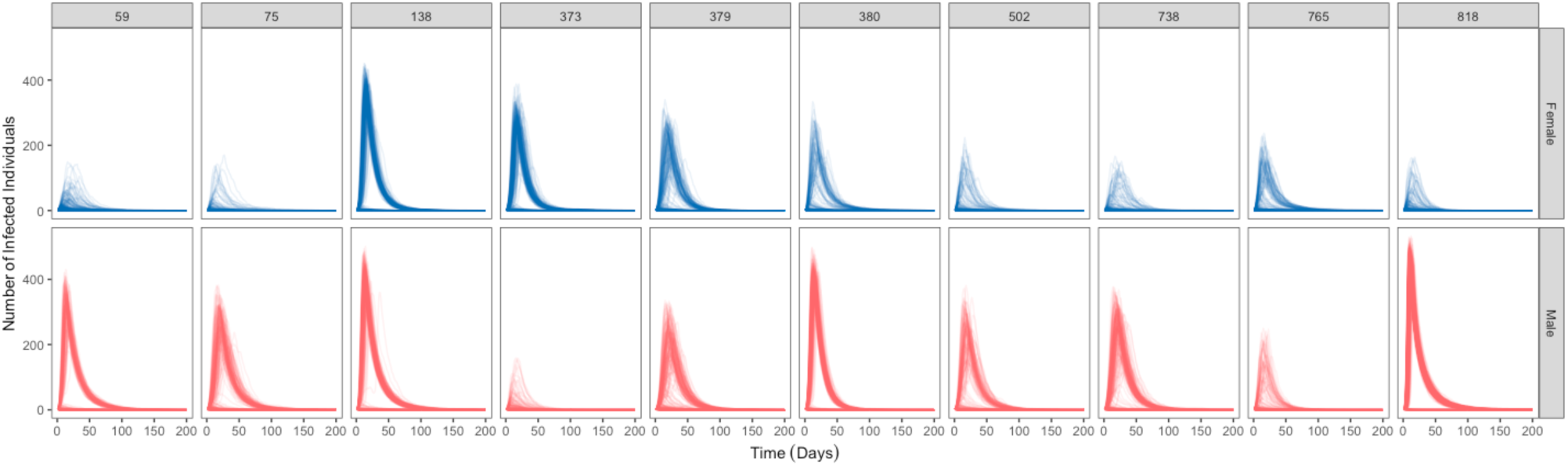
Simulation time courses of populations comprised of either male (red) or female (blue) individuals of the same and genetic background (columns) for simulation experiment #1. Across all of these simulations, parameters outside of host genetic background and sex are fixed; threshold radius (*r*) = 15mm, transmission efficiency (*τ*)=1 and scaled infectiousness (*η*)=2.

Our findings were robust to changes in the values of various combinations of sex, genetic background and variation in infectiousness, social aggregation and/or infection duration, alongside transmission efficiency, scaled infectiousness and threshold radius. Summary figures describing the fadeout likelihood, basic reproductive number (*R*_*0*_), maximum number of infected individuals, the time taken to reach the and maximum number of infected individuals, and outbreak duration, for every parameter combination are presented in Figures S1-12.

### Theoretical experiment #1

#### Individual variation in host infectiousness, social aggregation, and infection duration produced variation in population-level, pathogen transmission dynamics

The variation in empirical treatment groups produced distinct outbreaks of infectious disease in populations comprised solely of one genetic background and sex (Figures 1-2). Random forest analysis suggested that the two top predictors for outbreak likelihood were genetic and sex-specific variation (Figure 3a). Given a successful outbreak, host genetic and sex-specific variation also affected the maximum number of infected individuals at any given time step (Figures 2b & 3b) and outbreak duration (Figures 2c & 3c). However, host genetic background and sex were less important than the threshold radius used to derive social network degree distribution for both outcomes (Figure 3b & 3c) and less important than transmission efficiency for predicting the maximum number of infected individuals (Figure 3b).

**Figure 2.**
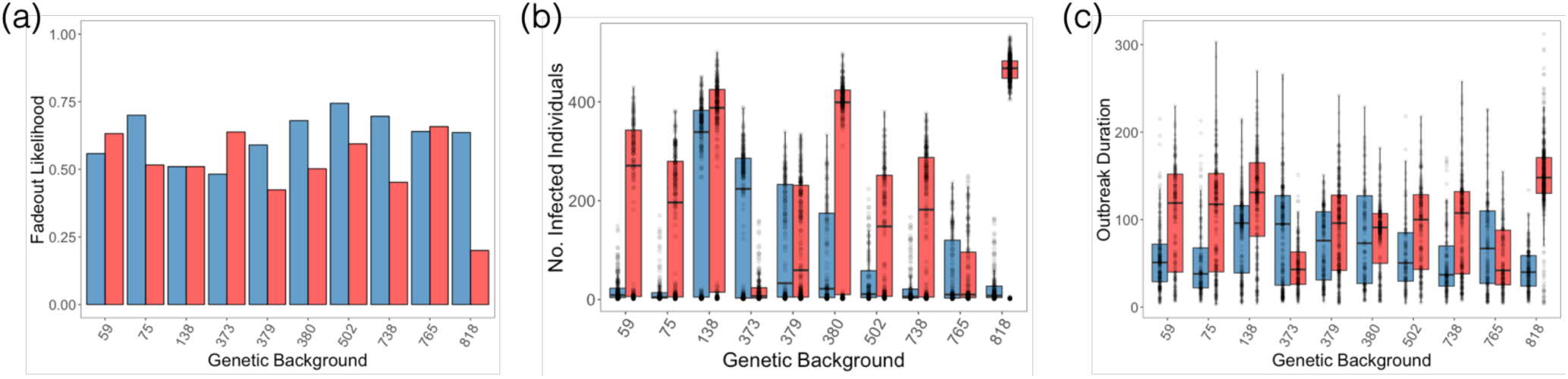
Summary statistics of time course simulations of populations comprised of either male (red) or female (blue) individuals of the same genetic background (x-axis). Across all of these simulations, parameters outside of host genetic background and sex are fixed; threshold radius (*r*) = 15mm, transmission efficiency (*τ*)=1 and scaled infectiousness (*η*)=2. All simulations were included to measure: (a) the proportion of simulations that resulted in fadeout; and, in the subset of simulations where fadeout did not occur, and disease spread from the index case; (b) the maximum number of infected individuals at any given time step; and (c) the number of time steps infected by the index case.

**Figure 3.**
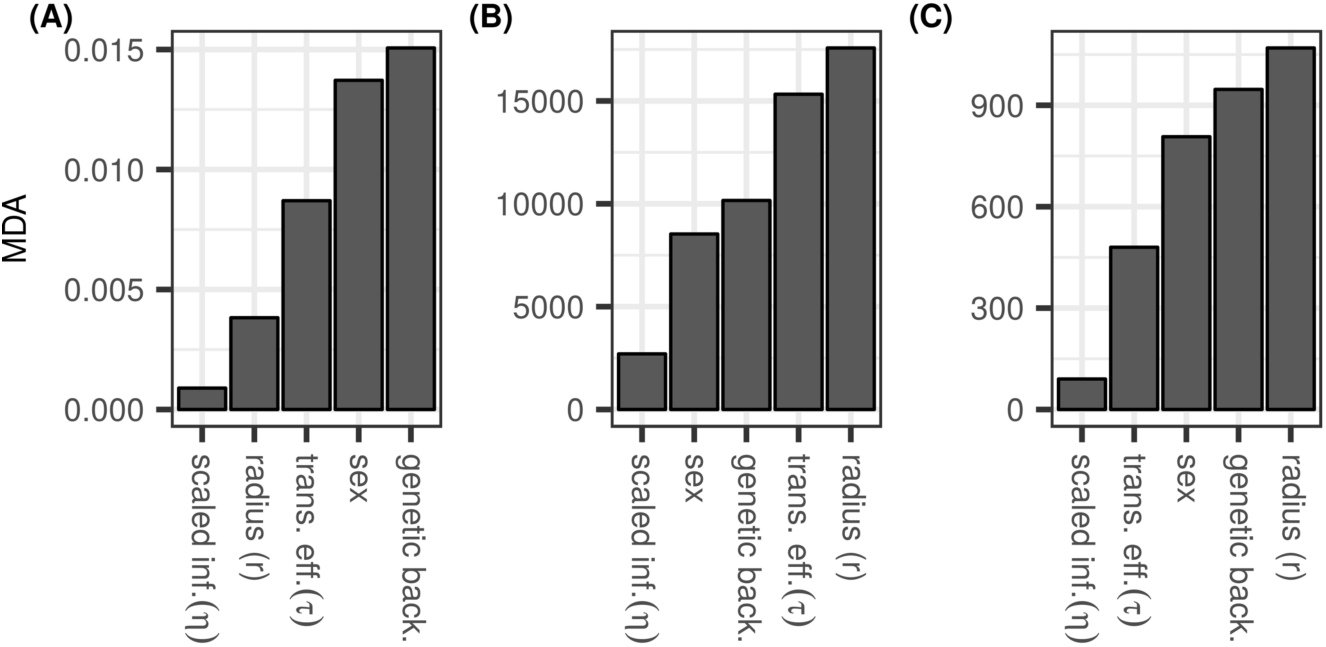
Results of variable importance analysis for theoretical experiment #1 with n=1000 trees using *cforest* function from the *party* package in *R*. Simulation variables are listed on x-axis. Y-axis describes variable importance (mean decrease in accuracy [MDA]). Which variables most determine: (A) whether the infection spread beyond initially infected individual? If so, which factors determine: (B) how many individuals infection reaches? and (C) how long it lasts?

### Theoretical experiment #2

#### Effects of the index case outweighed by heterogeneity in the susceptible population

The genetic background or sex of the index case did not alter outbreak dynamics in diverse populations where 20 empirical treatment groups were equally sampled to create a heterogeneous population (Figure 4 & 5). This was true for all outbreak descriptors (Figure 4 & 5). With no variation across empirical treatment groups, the importance of threshold radius, transmission efficiency, and scaled infectiousness influenced outbreaks, but in a consistent and predictable manner. Values conducive to greater infectiousness produced more likely and larger outbreaks (Figure 4 & 5). Based on the random forest analysis, threshold radius and transmission efficiency were the top two predictors for fadeout likelihood, maximum number of infected individuals, outbreak duration, and *R*_*0*_ (Figure S13).

**Figure 4.**
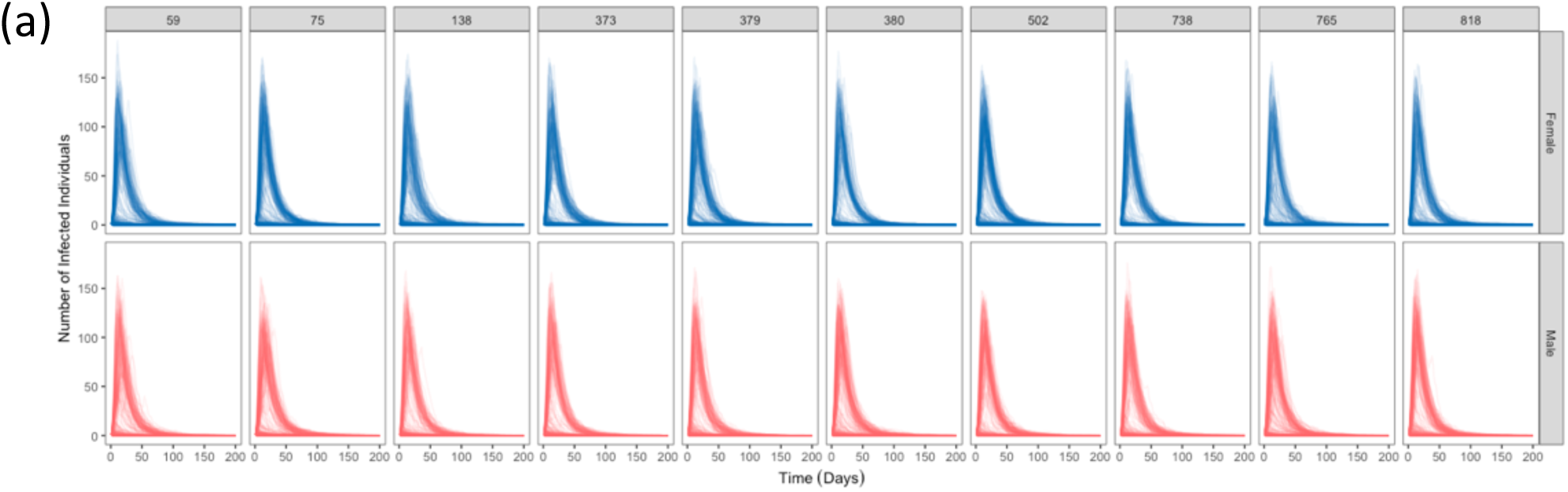
Simulation time courses of populations comprised of all ten genetic backgrounds and males (red), and females (blue) in equal proportion, where the index case of an outbreak is an individual of a specific genetic background and sex (simulation experiment #2). Across all of these simulations, other parameters are fixed: threshold radius (*r*) = 15mm, transmission efficiency (*τ*)=1 and scaled infectiousness (*η*)=2.

**Figure 5.**
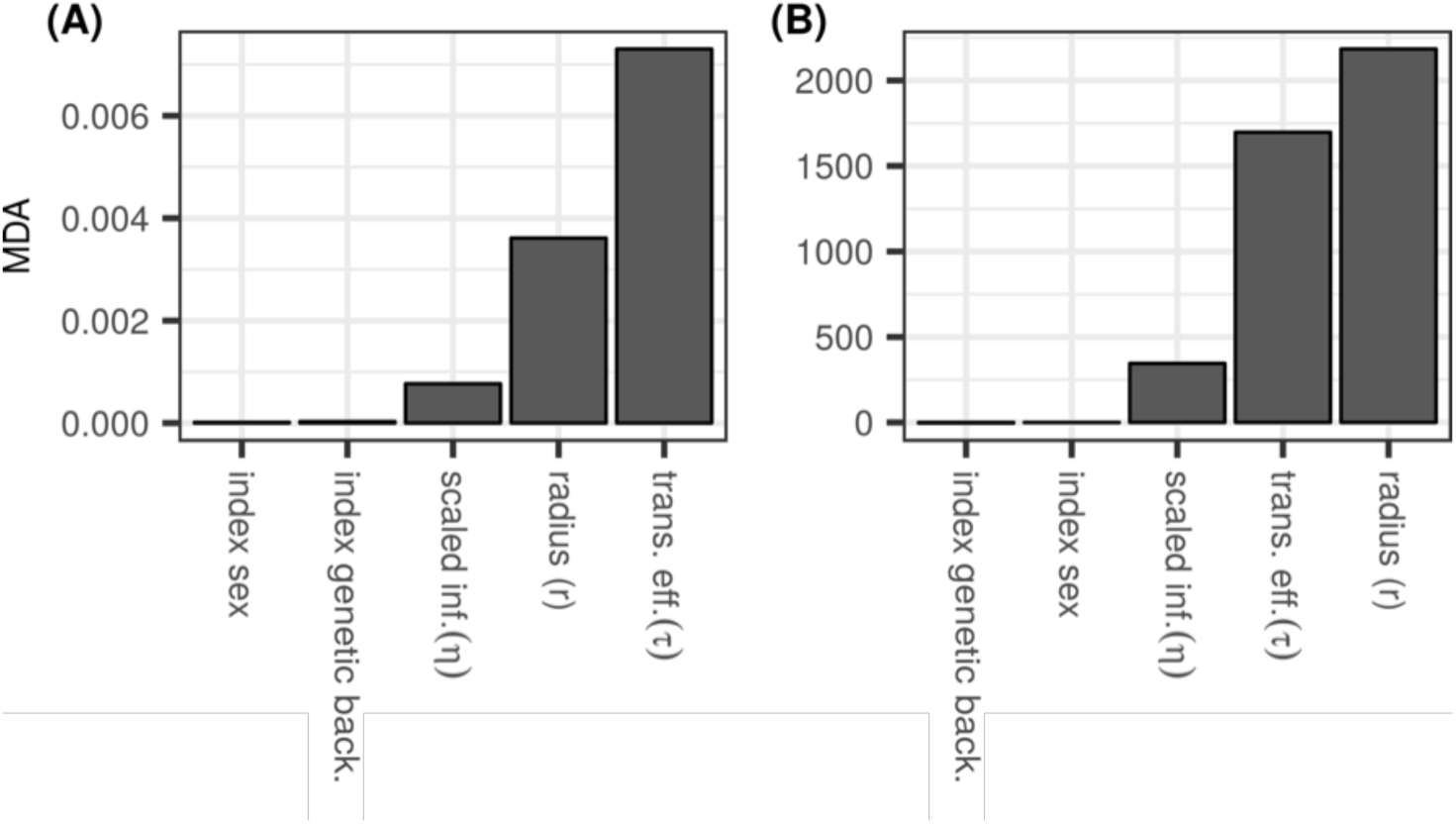
Results of variable importance analysis for Experiment #2 with n=1000 trees using *cforest* function from the *party* package in R. Variables are listed on x-axis. Y-axis describes variable importance (mean decrease in accuracy [MDA]). Which variables most determine: (A) whether the infection spread beyond initially infected individual? If so, which factors determine (B) how many individuals became infected?

### Theoretical experiment #3

#### Variation in infectiousness made outbreaks less likely to occur, spread to fewer individuals, and persist in the population for longer

Constraining the infectiousness of a population to the mean (0.23, 0.46 for scaled infectiousness (*η*) levels 1 and 2, respectively) of the empirical distribution increased the outbreak severity. This is clearly seen in outbreak time courses (Figure 6), making outbreaks are more likely (Figure 7a), infect more individuals (Figure 6 & 7b), and persist in the population for longer (Figure 6 & 7c). The only parameter that was not positively affected by constraining infectiousness to the mean was the time taken to reach the maximum number of infected individuals (Figure 7c). Here, limiting variation in infectiousness made outbreaks more predictable, reducing the variance of the time taken to reach the maximum number of infected individuals (Figure 7c). According to the random forest analysis, variation in infectiousness was the top predictor for whether or not an outbreak spread beyond the initially infected individual (Figure 8a).

**Figure 6.**
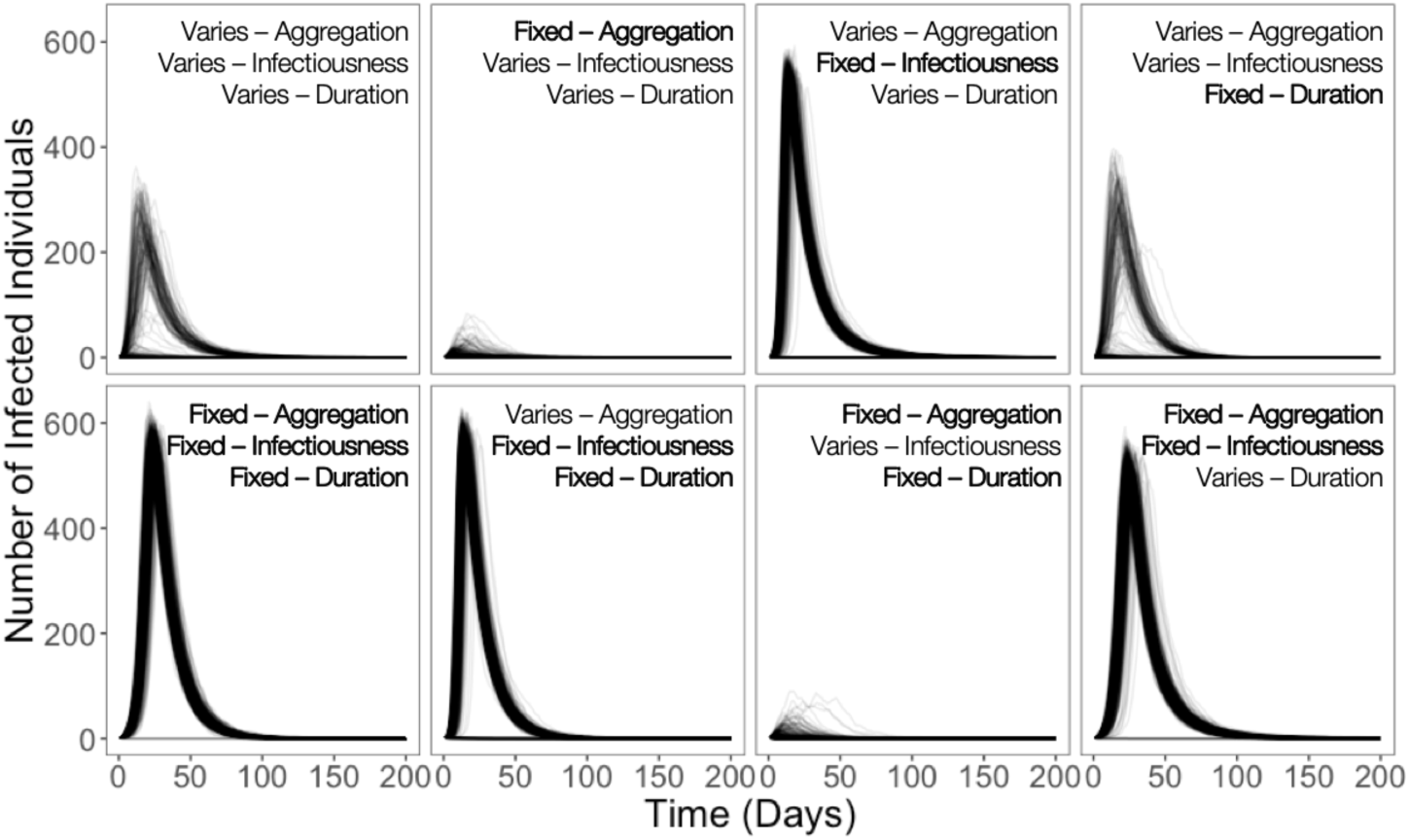
Simulation time courses of populations where aggregation, infectiousness and duration variation are derived from the breadth of the entire population’s variation rather than for a single genetic line and sex combination (simulation experiment #3). In each panel, the variation of a particular set of components is confined to the population’s mean. Across all of these simulations, parameters outside of host genetic background and sex are fixed: threshold radius = 15mm, transmission efficiency = 1 and scaled infectiousness = 2.

**Figure 7.**
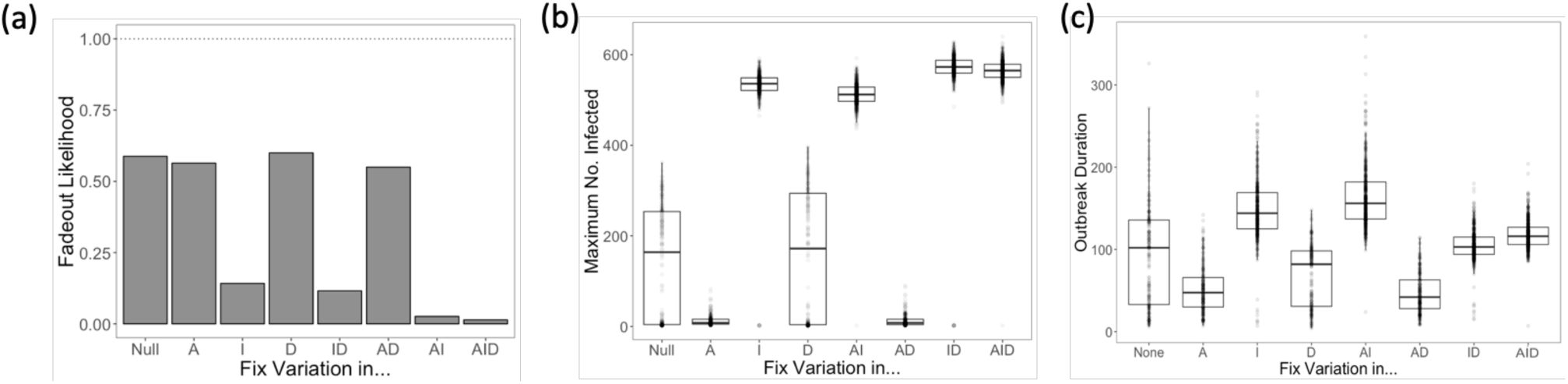
Summary statistics of time course simulations where individual variation is determined by the variation seen across all genetic backgrounds and sexes (simulation experiment #3). Across all of these simulations, threshold radius = 15mm, transmission efficiency = 1, and scaled infectiousness = 2. The x-axis of all panels sees variation in aggregation (A), infectiousness (I) and infection duration (D), and all their combinations fixed to the population mean. All simulations were included to measure: (a) the proportion of simulations that resulted in fadeout; (b) the maximum number of individuals infected during the simulation; (c) the time until maximum prevalence was reached.

**Figure 8.**
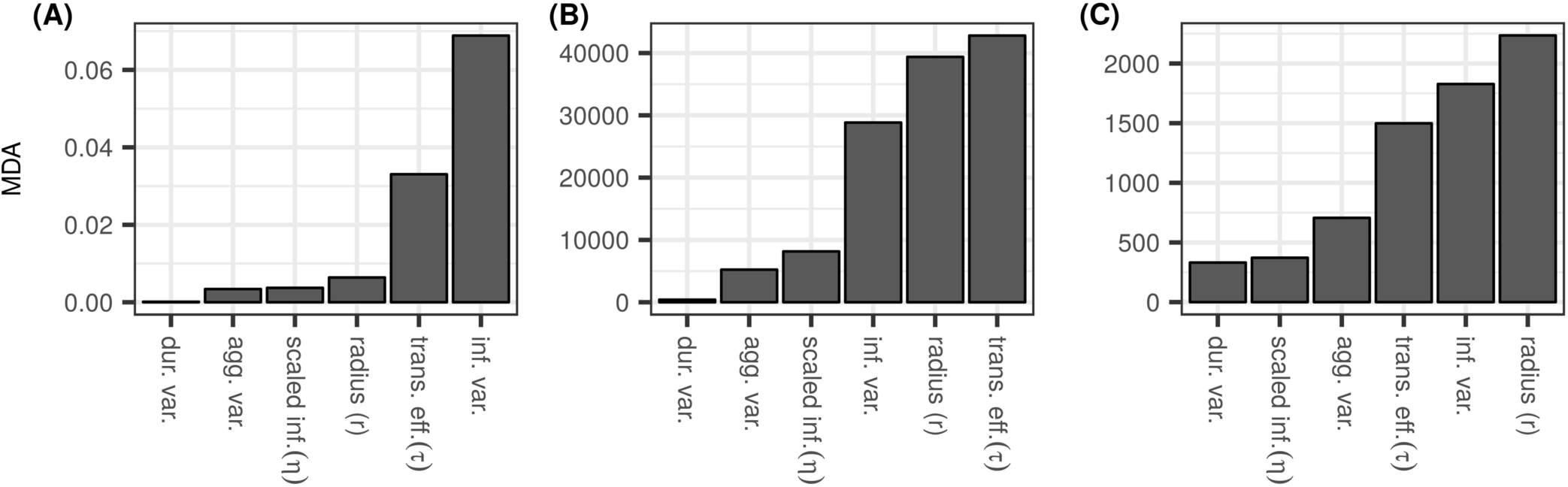
Results of variable importance analysis for Experiment 3 with n=1000 trees using *cforest* function from the *party* package in R. Variables are listed on x-axis. Y-axis describes variable importance (mean decrease in accuracy [MDA]). Which variables most determine: (A) whether the infection spread beyond initially infected individual? If so, which factors determine (B) how many individuals become infected and (C) how long it lasts.

#### Variation in social aggregation makes outbreaks more severe. It does not, however, influence outbreak likelihood

When social network degree distribution of simulated populations was confined to the mean of the empirical data (2, 3 and 4 for threshold radii of 10, 15 and 20mm respectively), outbreaks became less severe (Figure 6). Simulated DCV spread to fewer individuals (Figure 7b), at a slower rate (Figure 7c) and was quicker to die-out than in simulations where all three transmission components varied freely (Figure 7d).

#### Variation in disease-related mortality did not affect epidemic outcomes

When constrained to the mean of the empirical data (13.6 days), we found disease-related mortality had little to no effect on any aspect of disease outbreak (Figure 6). Constraining variation in disease-related mortality did not alter outbreak likelihood or severity (Figures 6 & 7a-c). This is supported by the random forest analysis which identified infection duration as the least important predictor across outbreak metrics (Figure 8a-c).

#### Variation in infectiousness, followed by social aggregation, is the most influential component of transmission

An increase in the maximum number of infected individuals is only seen when variation in infectiousness is constrained. Interestingly the same effect is seen in simulations where other traits are constrained alongside virus shedding, despite this differing substantially from the effects of social aggregation and infection duration when constrained alone (Figure 6-8). A similar, overruling effect is seen when social aggregation and infection duration are constrained simultaneously, and virus shedding varies freely; outbreak dynamics are similar to the cases where only aggregation is constrained (Figure 6-8).

#### Increasing the threshold radius increased outbreak severity but not likelihood

Manipulating the distance transmission can occur over, across all three theoretical experiments, made outbreaks more severe (Figure 9), but was not as strong of a predictor of outbreak likelihood (Figure 3, 5 & 8). Furthermore, we can see a non-linear relationship in the threshold radius when variation in social aggregation, virus shedding, and disease-related mortality were constrained to the population mean. When simulations derived social network according to one of three definitions of contact (10, 15 and 20mm), we see stark differences between 10 and 15mm, when compared to 15 and 20mm, indicating an important threshold value for the distance social interactions are drawn over. This distance was the most important predictor of outbreak parameters across almost all simulations (Figure 3, 5 & 8).

**Figure 9.**
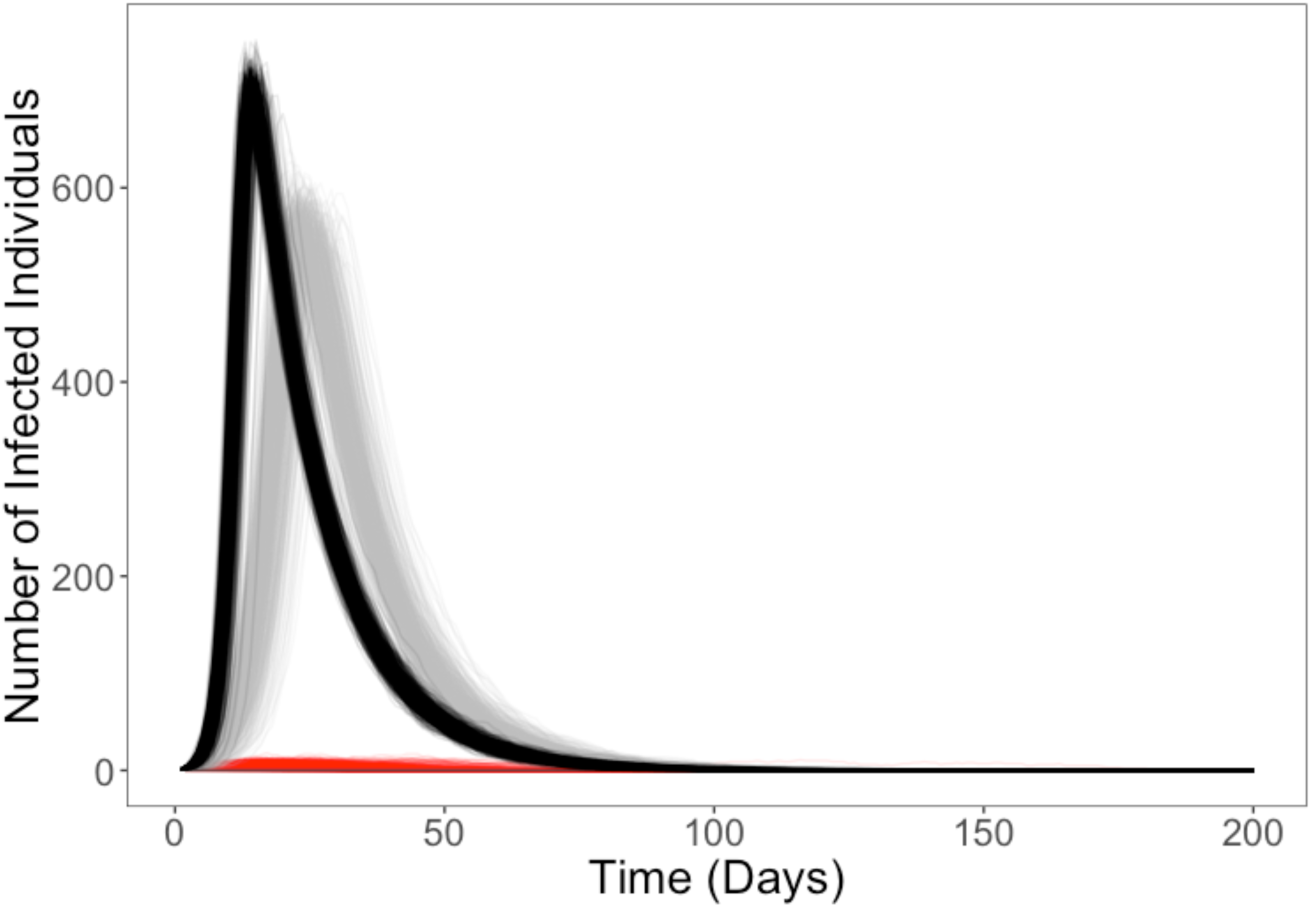
Simulation time courses of populations where aggregation, infectiousness and duration variation are fixed to the mean of the empirical population’s variation independently of genetic background and sex. A total of 3 radii were used to derive population social networks from empirically measured data, 10mm (red), 15mm (grey) and 20mm (black). Parameters outside of this infectious radius were constrained to one value; transmission efficiency = 1 and scaled infectiousness = 2.

## Discussion

We found substantial between-individual differences in disease transmission, constituting genetic and sex-specific variation in transmission potential. Crucially, in relatively homogenous populations comprised of single sex and genotype combinations, heterogeneity in the index case produced major differences in population-level outbreak dynamics, including making outbreaks more likely, broader reaching, and longer lasting. Despite the size of some of these differences however, variation in the index case’s transmission potential exerted little influence over population-level outbreak dynamics in diverse host populations. We also found that population-level variation in social aggregation, virus shedding, and disease-related mortality affected outbreak dynamics in starkly contrasting ways. This effect appeared to be linked to the population-level distribution of each respective host trait, with factors such as skewness and zero-inflation influencing how variation in each trait affected outbreak dynamics. Here, we discuss the traits of individuals that posed the greatest transmission risk and why they pose less risk in diverse susceptible populations. We also analyse the potential effects of “supersponges”, that represent no transmission risk to the susceptible population, and reflect on the broader implications of these results for mitigating the spread of disease in other host-pathogen systems.

To limit the pathogen transmission from high-risk individuals often requires expensive and continuous monitoring by experts. Focussing on classes of high-risk individuals is a more pragmatic approach to reducing the effect of heterogeneity in transmission potential, requiring less intensive monitoring protocols [6,40]. Additionally, as classes of individuals are identified using ranges of physiological or behavioural traits, classes are potentially more generalisable to other host-pathogen systems (e.g. sex, social dominance). In theoretical experiment 1, males from the RAL-818 genetic background were not only more likely to start an outbreak of infectious disease, but these outbreaks were also more severe than in other populations. This suggests these males represent a class of individuals with a high transmission risk. Interestingly, high-risk males are seen in a number of host-pathogen systems [41,42]. While high-risk male classes can be produced by a range of traits pertaining to sex-specific ecology or physiology, their occurrence across systems is likely driven by sexual selection shaping male traits affecting transmission [43]. For example, in the yellow-necked mouse, *Apodemus flavicollis*, males are thought to be a high-risk class due to a range of sex differences in their immune response, home range and contact rates [41]. Moreover, as male *Drosophila* exhibit a number of other traits with the potential to alter their transmission potential, such as male-male fighting [44], the transmission risk of RAL-818 males could increase further.

High-risk individuals, such as superspreaders, present a danger to current methods of disease control because they are capable of starting outbreaks of infectious disease that are difficult to predict and amplifying them once transmission begins [11,45]. This has driven pre-emptively identifying high-risk individuals to the forefront of epidemiology and disease ecology. However, in the second theoretical experiment we conducted, we found that starting outbreaks with individuals that differed in transmission potential, did not affect outbreak dynamics in diverse susceptible populations. The diversity of the susceptible population acted as an effective buffer to disease transmission, through low-competency individuals surrounding high-risk individuals. This effectively isolates high-risk individuals from the rest of the population. Our results suggest outbreaks are not solely driven by the traits of rare, high-risk individuals, but are also affected by the traits of the susceptible population. This finding overtly reflects a major strength of host diversity, increased protection from pathogens and parasites [35,46]. This finding bears a number of similarities with the broad observation that host diversity is central to the spread of disease [35,46,47]. The importance of host diversity to disease transmission is exemplified by the rapid spread of disease in crop monocultures due to crops being comprised of individuals with similar susceptibility [48–50]. A diverse host population exposes disease-causing agents to different environments that can apply unique selection pressures which prevent their proliferation. Similar transmission dynamics have also been observed in laboratory populations of the social spider, *Stegodyphus dumicola*, where transmission of a bacterial pathogen was affected by the boldness of the index case and the individuals it interacted with [11]. Together with our results, these findings do not suggest diversity in the susceptible population is a universal buffer to the effects of between-individual heterogeneity in disease transmission. Instead, this work highlights the necessity to characterise population diversity in the context of social interactions and networks as these may determine the relevance of this diversity. There are many traits across systems that bias social interactions, for example, such as sexual receptivity or personality type [51]. Should these traits bias contact between transmission classes, this may explain why social and contact networks rarely match.

Extreme phenotypes often play a key role in between-individual heterogeneity in disease transmission. However, being a relative term, ‘extreme’ phenotypes are defined by population-level variation. Constraining population-level variation in the amount of virus shed following infection to the population mean increased outbreak likelihood and severity. This was likely a result of the huge zero-inflation of the distribution of virus shedding, where many infected individuals did not shed virus. These individuals, previously termed ‘supersponges’ [52], represent the left-most extreme of the population distribution, and bore no transmission risk. ‘Supersponges’ present a potential explanation to the widely observed 20-80 rule across a range of host-pathogen and parasite systems, where, during an outbreak of infectious disease, 20% of hosts are responsible for 80% of transmission [53]. While some of the individuals that do not transmit infection may simply not get any transmission opportunity, others may be supersponges and therefore incapable of transmitting disease.

An important caveat of our results is that because we did not measure social aggregation, virus shedding and lifespan simultaneously we cannot account for how they might covary in individuals. We therefore allow them to co-occur in hosts randomly, which may contradict associations produced in nature or predicted by hypotheses [54,55]. This is particularly true for how we derived contact between susceptible and infected individuals from empirical data on social aggregation. An important caveat of contact in our simulations is that it is derived from social aggregation arenas containing 10-12 flies and measuring 55mm wide. We considered populations of 1000 individuals whose social network was derived by scaling-up the aggregation of considerably smaller populations. This approach was necessitated by the experimental demands of measuring social aggregation [56]. However, it would be pertinent to experimentally test how *Drosophila* social aggregation changes with population size, particularly as this is a common issue where empirical data is integrated with *in silico* work. Moreover, larger populations often have different structures and more modular social networks which have been shown to facilitate or prevent the spread of disease [57,58]. As our social aggregation data comes from Petri dishes containing only males or females from a single genetic background, we cannot account for how aggregation might change in more diverse and larger populations. Additionally, as very little is known about DCV transmission, the distance DCV transmission was able to occur over was inherently arbitrary, despite this issue being accounted for by threshold radius. Understanding how distance affects pathogen transmission or definitions of what constitutes a contact is a hugely influential relationship which is poorly described in many host-pathogen systems (White et al. 2018). The high levels of control in laboratory populations of *Drosophila* offer an ideal system to characterise this relationship.

Our work bears a number of consequences for understanding how between-individual heterogeneity in disease transmission is determined and how it could affect outbreak dynamics. We show that variation in key traits in individuals can dramatically affect population-level transmission, surmounting to genetic and sex-specific variation in transmission potential. Importantly, the influence of this variation is dramatically affected by susceptible population diversity and the distribution of population-level variation. These results support the observations of other systems that suggest the traits of susceptible individuals can exert significant influence over transmission. This is particularly relevant to populations with low genetic diversity, such as agricultural monocultures, as this lack of diversity increases the risk of explosive outbreaks [49,50,59]. This result also applies to organisms with structured populations, such as familial herds, or nests, as this structure could produce localised groups of high-risk individuals. Our work posits the merits of integrating data collected in highly controlled laboratory experiments with simulations capable of extrapolating this information to larger populations. Simulation experiments may also help generate hypotheses of particular interest in a laboratory setting [60]. Given that there are many other protocols available to the *D. melanogaster* model system, as well as many other systems, our holistic approach provides an ideal tool to furthering our understanding of how rare, high-risk individuals affect population-level transmission dynamics.

## Funding

J.A.S-J was funded by a NERC E3 DTP PhD studentship awarded to the University of Edinburgh. L.A.W. was funded by the National Science Foundation (GRFP-00039202 and DEB-1701069), the University of Minnesota Informatics Institute, and the National Socio-Environmental Synthesis Center (SESYNC) under funding received from the National Science Foundation DBI-1639145. J.A.S-J and L.A.W. received funding for a research exchange from the Infectious Disease Evolution Across Scales RCN funded by NSF.M.E.C. was funded by National Science Foundation (DEB-1413925 and 1654609) and CVM Research Office UMN Ag Experiment Station General Ag Research Funds. The authors acknowledge the Minnesota Supercomputing Institute (MSI) at the University of Minnesota for providing resources that contributed to the research results reported within this paper. URL: http://www.msi.umn.edu. P.F.V was supported by a Branco Weiss fellowship (https://brancoweissfellowship.org/) and a Chancellor’s Fellowship (School of Biological Sciences, University of Edinburgh).

## Notes

https://github.com/whit1951/Drosophila

## References

1. Craft ME. Infectious disease transmission and contact networks in wildlife and livestock. Philos Trans R Soc Lond B Biol Sci. 2015;370: 20140107-. doi:10.1098/rstb.2014.0107

2. Gopinath S, Lichtman JS, Bouley DM, Elias JE, Monack DM. Role of disease-associated tolerance in infectious superspreaders. Proc Natl Acad Sci U S A. 2014;111: 15780–15785. doi:10.1073/pnas.1409968111

3. Lloyd-Smith JO, Schreiber SJ, Kopp PE, Getz WM, Schreiber SJ, Kopp PE, et al. Superspreading and the effect of individual variation on disease emergence. Nature. 2005;438: 355–9. doi:10.1038/nature04153

4. White LA, Forester JD, Craft ME. Using contact networks to explore mechanisms of parasite transmission in wildlife. Biol Rev. 2017;92: 389–409. doi:10.1111/brv.12236

5. Anderson RM, May RM. The Population Dynamics of Microparasites and Their Invertebrate Hosts. Philos Trans R Soc B Biol Sci. 1981;291: 451–524. doi:10.1098/rstb.1981.0005

6. VanderWaal KL, Ezenwa VO. Heterogeneity in pathogen transmission: mechanisms and methodology. Funct Ecol. 2016;30: 1606–1622. doi:10.1111/1365-2435.12645

7. Hawley DM, Altizer SM. Disease ecology meets ecological immunology: understanding the links between organismal immunity and infection dynamics in natural populations. Funct Ecol. 2011;25: 48–60. doi:10.1111/j.1365-2435.2010.01753.x

8. Lopes PC, Block P, König B. Infection-induced behavioural changes reduce connectivity and the potential for disease spread in wild mice contact networks. Sci Rep. 2016;6: 31790. doi:10.1038/srep31790

9. Susi H, Vale PF, Laine A-L. Host Genotype and Coinfection Modify the Relationship of within and between Host Transmission. Am Nat. 2015;186: 252–263. doi:10.1086/682069

10. Vale Pedro F., Choisy Marc, Little Tom J. Host nutrition alters the variance in parasite transmission potential. Biol Lett. 2013;9: 20121145. doi:10.1098/rsbl.2012.1145

11. Keiser CN, Pinter-Wollman N, Ziemba MJ, Kothamasu KS, Pruitt JN. The index case is not enough: Variation among individuals, groups and social networks modify bacterial transmission dynamics. J Anim Ecol. 2017; 1–10. doi:10.1111/1365-2656.12729

12. Lloyd-Smith James O., Getz Wayne M., Westerhoff Hans V. Frequency–dependent incidence in models of sexually transmitted diseases: portrayal of pair–based transmission and effects of illness on contact behaviour. Proc R Soc Lond B Biol Sci. 2004;271: 625–634. doi:10.1098/rspb.2003.2632

13. May RM, Anderson RM. Transmission dynamics of HIV infection. Nature. 1987;326: 137–42. doi:10.1038/326137a0

14. White LA, Forester JD, Craft ME. Covariation between the physiological and behavioral components of pathogen transmission: host heterogeneity determines epidemic outcomes. Oikos. 2018;127: 538–552. doi:10.1111/oik.04527

15. Elderd BD, Keeling M, Woolhouse M, May R, Davies G, Grenfell B, et al. Developing Models of Disease Transmission: Insights from Ecological Studies of Insects and Their Baculoviruses. Heitman J, editor. PLoS Pathog. 2013;9: e1003372–e1003372. doi:10.1371/journal.ppat.1003372

16. Gandon S, Day T, Metcalf CJE, Grenfell BT. Forecasting Epidemiological and Evolutionary Dynamics of Infectious Diseases. Trends Ecol Evol. 2016;31: 776–788. doi:10.1016/j.tree.2016.07.010

17. Cross PC, Johnson PLF, Lloyd-Smith JO, Getz WM. Utility of R0 as a predictor of disease invasion in structured populations. J R Soc Interface. 2007;4: 315–324. doi:10.1098/rsif.2006.0185

18. Lloyd-Smith JO, Schreiber SJ, Getz WM. Moving beyond averages: individual-level variation in disease transmission. In: Gumel AB, Castillo-Chavez C, Mickens RE, Clemence DP, editors. Contemporary Mathematics. Providence, Rhode Island: American Mathematical Society; 2006. pp. 235–258. doi:10.1090/conm/410/07730

19. Lloyd-Smith JO, Schreiber S, Kopp P, Getz W. Superspreading and the effect of individual variation on disease emergence. Nature. 2005;438: 355–359. doi:10.1038/nature04153

20. Keiser CN, Pinter-Wollman N, Augustine DA, Ziemba MJ, Hao L, Lawrence JG, et al. Individual differences in boldness influence patterns of social interactions and the transmission of cuticular bacteria among group-mates. Proc R Soc B Biol Sci. 2016;283: 20160457–20160457. doi:10.1098/rspb.2016.0457

21. Susi H, Barrès B, Vale PF, Laine A-L. Co-infection alters population dynamics of infectious disease. Nat Commun. 2015;6. doi:10.1038/ncomms6975

22. Lloyd-Smith JO, George D, Pepin KM, Pitzer VE, Pulliam JRC, Dobson AP, et al. Epidemic dynamics at the human-animal interface. Science. 2009;326: 1362–1367. doi:10.1126/science.1177345

23. McCallum H, Fenton A, Hudson PJ, Lee B, Levick B, Norman R, et al. Breaking beta: deconstructing the parasite transmission function. Philos Trans B. 2017;372: 20160084. doi:10.1098/rstb.2016.0084

24. Siva-Jothy JA, Vale PF. Viral infection causes sex-specific changes in fruit fly social aggregation behaviour. bioRxiv. 2019; 630913. doi:10.1101/630913

25. Siva-Jothy JA, Vale PF. Dissecting genetic and sex-specific host heterogeneity in pathogen transmission potential. bioRxiv. 2019; 733915. doi:10.1101/733915

26. Keebaugh ES, Schlenke TA. Insights from natural host–parasite interactions: The Drosophila model. Dev Comp Immunol. 2014;42: 111–123. doi:10.1016/j.dci.2013.06.001

27. Mackay TFC, Richards S, Stone EA, Barbadilla A, Ayroles JF, Zhu D, et al. The Drosophila melanogaster Genetic Reference Panel. Nature. 2012;482: 173–8. doi:10.1038/nature10811

28. Magwire MM, Fabian DK, Schweyen H, Cao C, Longdon B, Bayer F, et al. Genome-Wide Association Studies Reveal a Simple Genetic Basis of Resistance to Naturally Coevolving Viruses in Drosophila melanogaster. PLoS Genet. 2012;8. doi:10.1371/journal.pgen.1003057

29. Siva-Jothy JA, Monteith KM, Vale PF. Navigating infection risk during oviposition and cannibalistic foraging in a holometabolous insect. Behav Ecol. 2018; doi:10.1093/beheco/ary106

30. Merkling SH, van Rij RP. Analysis of resistance and tolerance to virus infection in Drosophila. Nat Protoc. 2015;10: 1084–97. doi:10.1038/nprot.2015.071

31. Anderson R, May R. Infectious diseases of humans. Oxford: Oxford University Press; 1991.

32. Huszar T, Imler J. Drosophila Viruses and the Study of Antiviral Host-Defense. Advances in Virus Research. Academic Press; 2008. pp. 227–265.

33. Webster CL, Waldron FM, Robertson S, Crowson D, Ferrari G, Quintana JF, et al. The Discovery, Distribution, and Evolution of Viruses Associated with Drosophila melanogaster. PLoS Biol. 2015;13: e1002210–e1002210. doi:10.1371/journal.pbio.1002210

34. Kapun M, Nolte V, Flatt T, Schlötterer C. Host Range and Specificity of the Drosophila C Virus. PLoS ONE. 2010;5. doi:10.1371/journal.pone.0012421

35. Ostfeld RS, Keesing F. Effects of Host Diversity on Infectious Disease. Annu Rev Ecol Evol Syst. 2012;43: 157–182. doi:10.1146/annurev-ecolsys-102710-145022

36. Cutler DR, Edwards TC, Beard KH, Cutler A, Hess KT, Gibson J, et al. Random forests for classification in ecology. Ecology. 2007;88: 2783–2792. doi:10.1890/07-0539.1

37. Breiman L. Random forests. Mach Learn. 2001;45: 5–32. doi:10.1023/A:1010933404324

38. Strobl C, Hothorn T, Zeileis A. Party on! A new, conditional variable-importance measure for random forests available in the party package. R J. 2009;1: 14–17.

39. Strobl C, Boulesteix A-L, Zeileis A, Hothorn T. Bias in random forest variable importance measures: illustrations, sources and a solution. BMC Bioinformatics. 2007;8: 25. doi:10.1186/1471-2105-8-25

40. Drewe JA. Who infects whom? Social networks and tuberculosis transmission in wild meerkats. Proc Biol Sci. 2009;277: 633–642. doi:10.1098/rspb.2009.1775

41. Ferrari N, Cattadori IM, Nespereira J, Rizzoli A, Hudson PJ. The role of host sex in parasite dynamics: field experiments on the yellow-necked mouse Apodemus flavicollis. Ecol Lett. 2004;7: 88–94. doi:10.1046/j.1461-0248.2003.00552.x

42. Grear DA, Perkins SE, Hudson PJ. Does elevated testosterone result in increased exposure and transmission of parasites? Ecol Lett. 2009;12: 528–537. doi:10.1111/j.1461-0248.2009.01306.x

43. Zuk M, McKean KA. Sex differences in parasite infections: Patterns and processes. Int J Parasitol. 1996;26: 1009–1024. doi:10.1016/S0020-7519(96)80001-4

44. Baxter CM, Barnett R, Dukas R. Aggression, mate guarding and fitness in male fruit flies. Anim Behav. 2015;109: 235–241. doi:10.1016/j.anbehav.2015.08.023

45. Craft ME, Caillaud D. Network Models: An Underutilized Tool in Wildlife Epidemiology? In: Interdisciplinary Perspectives on Infectious Diseases [Internet]. 2011 [cited 1 Jun 2019]. doi:10.1155/2011/676949

46. Lively CM. The Effect of Host Genetic Diversity on Disease Spread. Am Nat. 2010;175: E149–E152. doi:10.1086/652430

47. Altermatt F, Ebert D. Genetic diversity of Daphnia magna populations enhances resistance to parasites. Ecol Lett. 2008;11: 918–928. doi:10.1111/j.1461-0248.2008.01203.x

48. Mitchell CE, Tilman D, Groth JV. Effects of Grassland Plant Species Diversity, Abundance, and Composition on Foliar Fungal Disease. Ecology. 2002;83: 1713–1726. doi:10.1890/0012-9658(2002)083[1713:EOGPSD]2.0.CO;2

49. Pilet F, Chacón G, Forbes GA, Andrivon D. Protection of Susceptible Potato Cultivars Against Late Blight in Mixtures Increases with Decreasing Disease Pressure. Phytopathology. 2006;96: 777–783. doi:10.1094/PHYTO-96-0777

50. Zhu Y, Chen H, Fan J, Wang Y, Li Y, Chen J, et al. Genetic diversity and disease control in rice. Nature. 2000;406: 718. doi:10.1038/35021046

51. Keiser CN, Howell KA, Pinter-Wollman N, Pruitt JN. Personality composition alters the transmission of cuticular bacteria in social groups. Biol Lett. 2016;12: 20160297. doi:10.1098/rsbl.2016.0297

52. Barron DG, Gervasi SS, Pruitt JN, Martin LB. Behavioral competence: How host behaviors can interact to influence parasite transmission risk. Curr Opin Behav Sci. 2015;6: 35–40. doi:10.1016/j.cobeha.2015.08.002

53. Woolhouse MEJ, Dye C, Etard J-F, Smith T, Charlwood JD, Garnett GP, et al. Heterogeneities in the transmission of infectious agents?: Proc Natl Acad Sci U S A. 1997;94: 338–342.

54. Réale D, Garant D, Humphries MM, Bergeron P, Careau V, Montiglio P-O. Personality and the emergence of the pace-of-life syndrome concept at the population level. Philos Trans R Soc B Biol Sci. 2010;365: 4051–4063. doi:10.1098/rstb.2010.0208

55. Vazquez-Prokopec GM, Perkins TA, Waller LA, Lloyd AL, Reiner RC, Scott TW, et al. Coupled Heterogeneities and Their Impact on Parasite Transmission and Control. Trends Parasitol. 2016;32: 356–367. doi:10.1016/j.pt.2016.01.001

56. Simon AF, Chou M-T, Salazar ED, Nicholson T, Saini N, Metchev S, et al. A simple assay to study social behavior in Drosophila: measurement of social space within a group. Genes Brain Behav. 2012;11: 243–52. doi:10.1111/j.1601-183X.2011.00740.x

57. Nunn CL, Jordán F, McCabe CM, Verdolin JL, Fewell JH. Infectious disease and group size: more than just a numbers game. Philos Trans R Soc Lond B Biol Sci. 2015;370.

58. Sah P, Leu ST, Cross PC, Hudson PJ, Bansal S. Unraveling the disease consequences and mechanisms of modular structure in animal social networks. Proc Natl Acad Sci. 2017;114: 4165–4170. doi:10.1073/pnas.1613616114

59. Wallace R, Wallace RG. Blowback: new formal perspectives on agriculturally driven pathogen evolution and spread. Epidemiol Infect. 2015;143: 2068–2080. doi:10.1017/S0950268814000077

60. Restif O, Hayman DTS, Pulliam JRC, Plowright RK, George DB, Luis AD, et al. Model-guided fieldwork: practical guidelines for multidisciplinary research on wildlife ecological and epidemiological dynamics. Ecol Lett. 2012;15: 1083–1094. doi:10.1111/j.1461-0248.2012.01836.x

