## Supplementary material for "Population-Level Disease Dynamics Reflect Individual Heterogeneities in Transmission": Figures S1-12

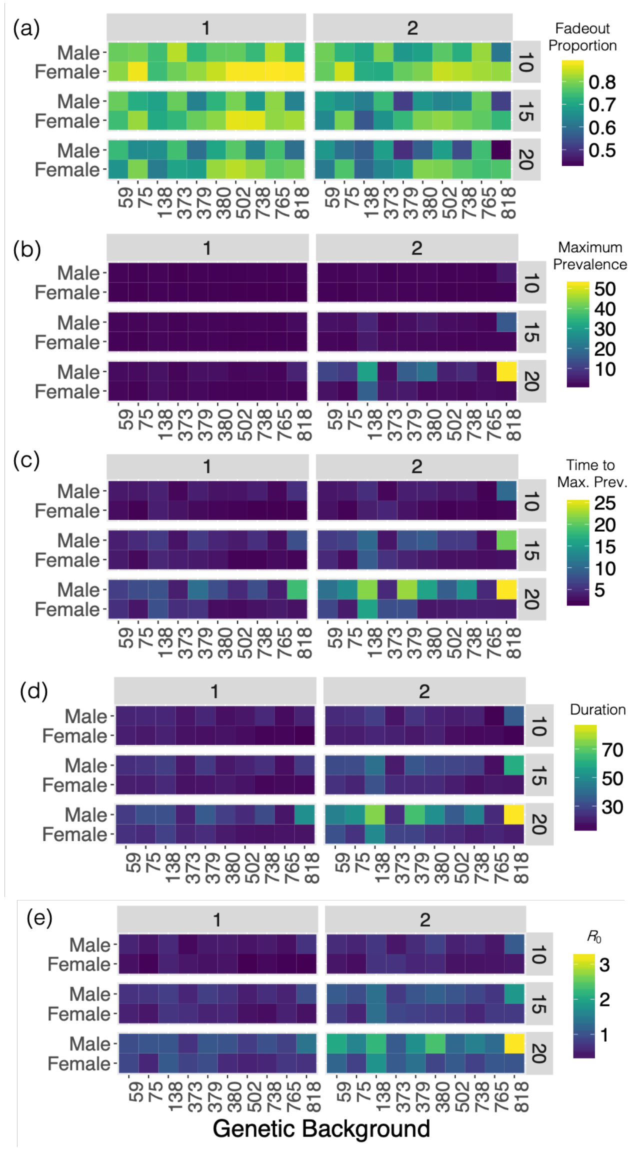


**Figure S1. Theoretical experiment #1 (τ=0.1)** - Mean (a) fadeout proportion, (b) maximum number of infected individuals (maximum prevalence), (c) the number of time steps taken to reach the maximum number of infected individuals (time to Max. Prev.) (d) the number of time steps outbreaks lasted for (duration) and (e) the number of secondary infections caused by the first infected individual ($R_0$) of outbreaks simulated in theoretical populations in experiment #1. Theoretical populations were comprised of individuals of the same sex and genetic background. Across these simulations, pathogen transmission efficiency (τ) was 0.1, infectiousness (η) was scaled by 1 or 2 (x-axis facets) and contact network connections were based on flies aggregating within 10, 15 or 20mm of one another (r; y-axis facets).

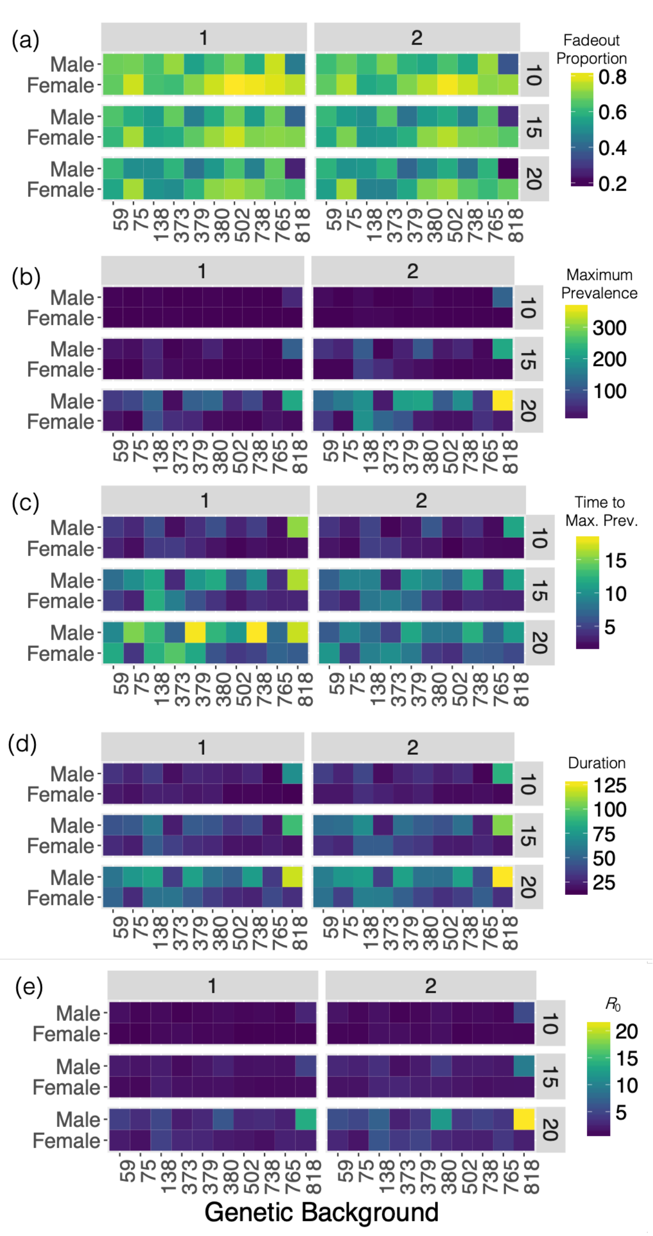


**Figure S2. Theoretical experiment #1 (τ=0.5)** - Mean (a) fadeout proportion, (b) maximum number of infected individuals (maximum prevalence), (c) the number of time steps taken to reach the maximum number of infected individuals (time to Max. Prev.) (d) the number of time steps outbreaks lasted for (duration) and (e) the number of secondary infections caused by the first infected individual ($R_0$) of outbreaks simulated in theoretical populations in experiment #1. Theoretical populations were comprised of individuals of the same sex and genetic background. Across these simulations, pathogen transmission efficiency (τ) was 0.1, infectiousness (η) was scaled by 1 or 2 (x-axis facets) and contact network connections were based on flies aggregating within 10, 15 or 20mm of one another (r; y-axis facets).

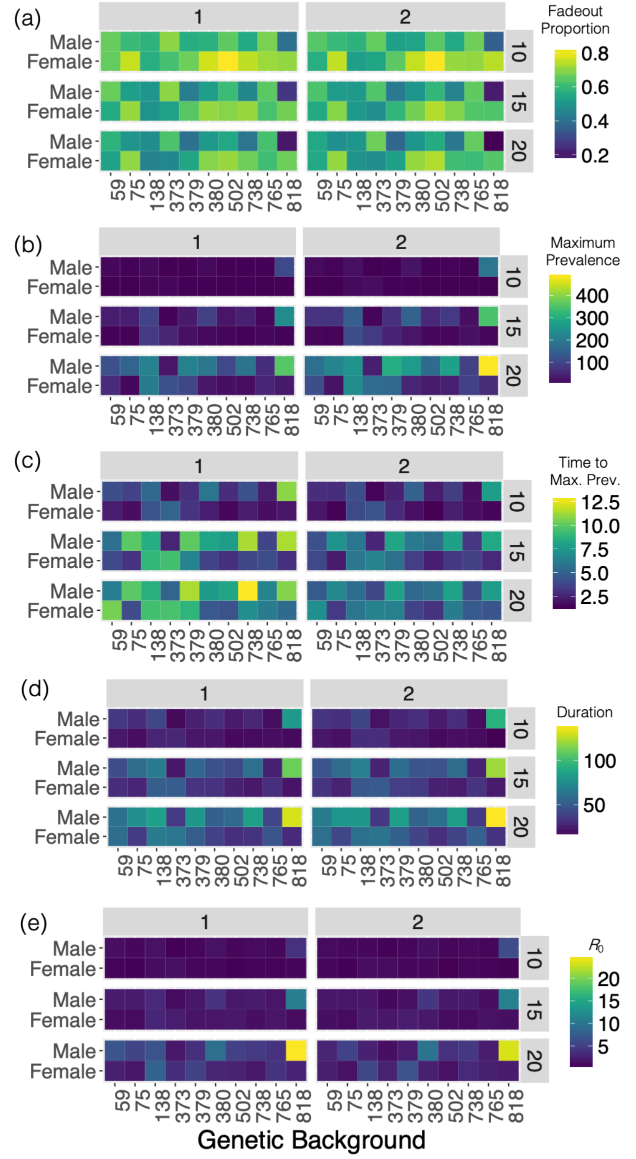


**Figure S3. Theoretical experiment #1 (τ=1.0)** - Mean (a) fadeout proportion, (b) maximum number of infected individuals (maximum prevalence), (c) the number of time steps taken to reach the maximum number of infected individuals (time to Max. Prev.) (d) the number of time steps outbreaks lasted for (duration) and (e) the number of secondary infections caused by the first infected individual ($R_0$) of outbreaks simulated in theoretical populations in experiment #1. Theoretical populations were comprised of individuals of the same sex and genetic background. Across these simulations, pathogen transmission efficiency (τ) was 0.1, infectiousness (η) was scaled by 1 or 2 (x-axis facets) and contact network connections were based on flies aggregating within 10, 15 or 20mm of one another (r; y-axis facets).

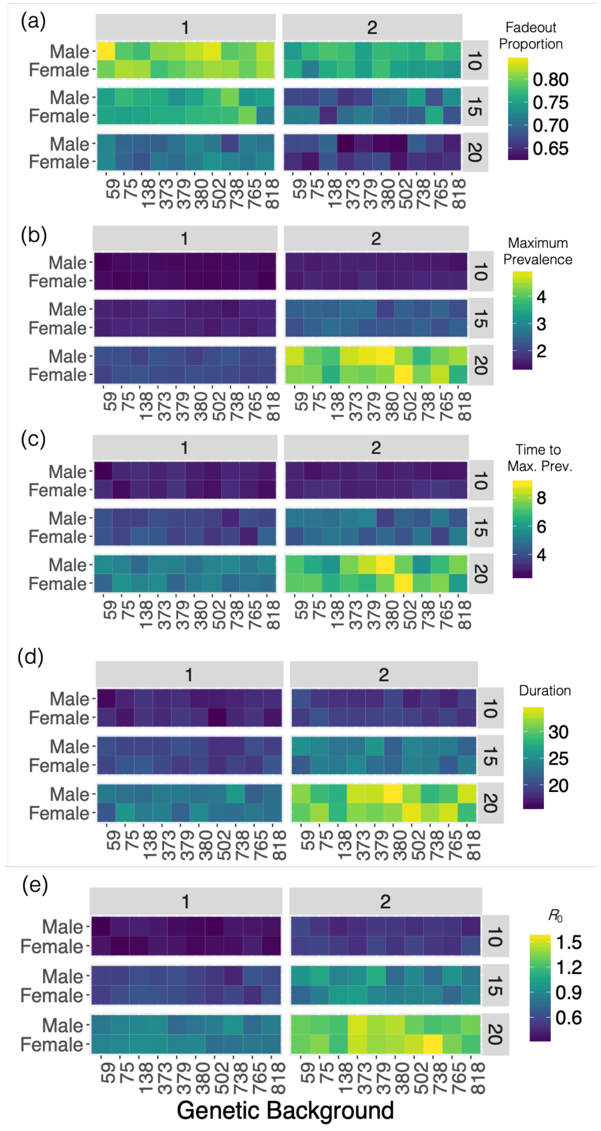


**Figure S4. Theoretical experiment #2 (τ=0.1)** - Mean (a) fadeout proportion, (b) maximum number of infected individuals (maximum prevalence), (c) the number of time steps taken to reach the maximum number of infected individuals (time to Max. Prev.) (d) the number of time steps outbreaks lasted for (duration) and (e) the number of secondary infections caused by the first infected individual ($R_0$) of outbreaks simulated in theoretical populations in experiment #2. Theoretical populations were comprised of 50 individuals from each combination of sex and genetic background. Across these simulations, pathogen transmission efficiency (τ) was 0.1, infectiousness (η) was scaled by 1 or 2 (x-axis facets) and contact network connections were based on flies aggregating within 10, 15 or 20mm of one another (r; y-axis facets).

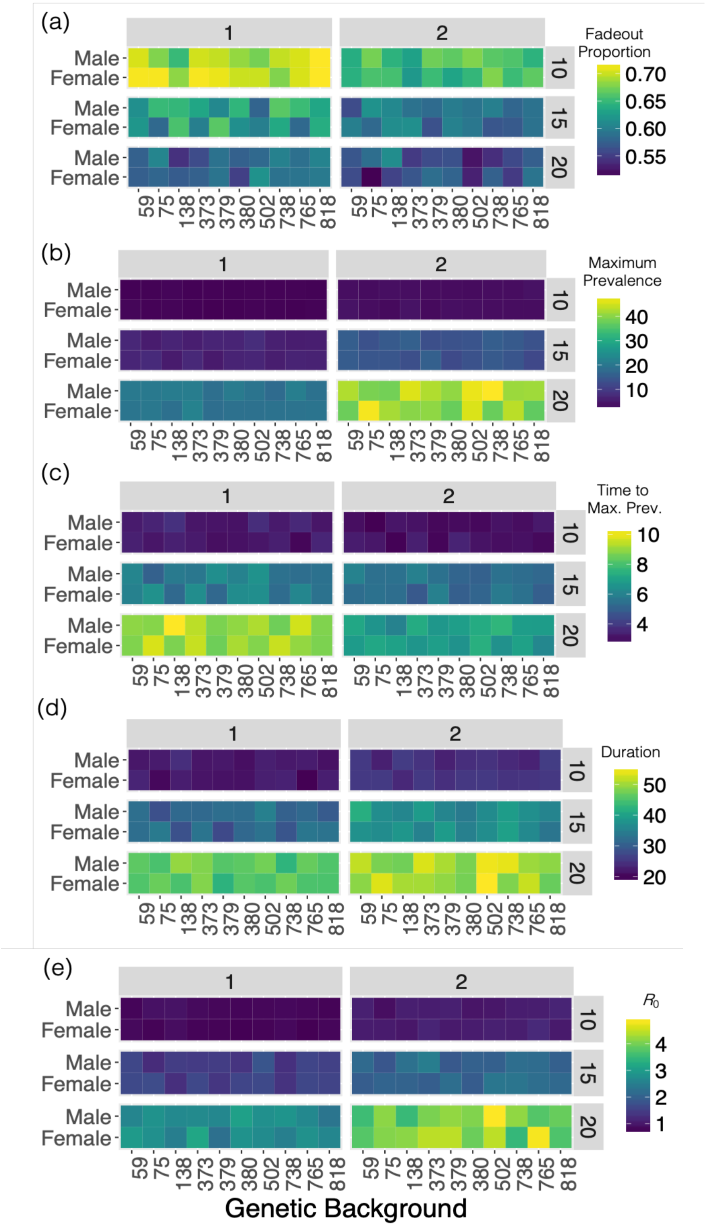


**Figure S5. Theoretical experiment #2 (τ=0.5)** - Mean (a) fadeout proportion, (b) maximum number of infected individuals (maximum prevalence), (c) the number of time steps taken to reach the maximum number of infected individuals (time to Max. Prev.) (d) the number of time steps outbreaks lasted for (duration) and (e) the number of secondary infections caused by the first infected individual ($R_0$) of outbreaks simulated in theoretical populations in experiment #2. Theoretical populations were comprised of 50 individuals from each combination of sex and genetic background. Across these simulations, pathogen transmission efficiency (τ) was 0.5, infectiousness (η) was scaled by 1 or 2 (x-axis facets) and contact network connections were based on flies aggregating within 10, 15 or 20mm of one another (r; y-axis facets).

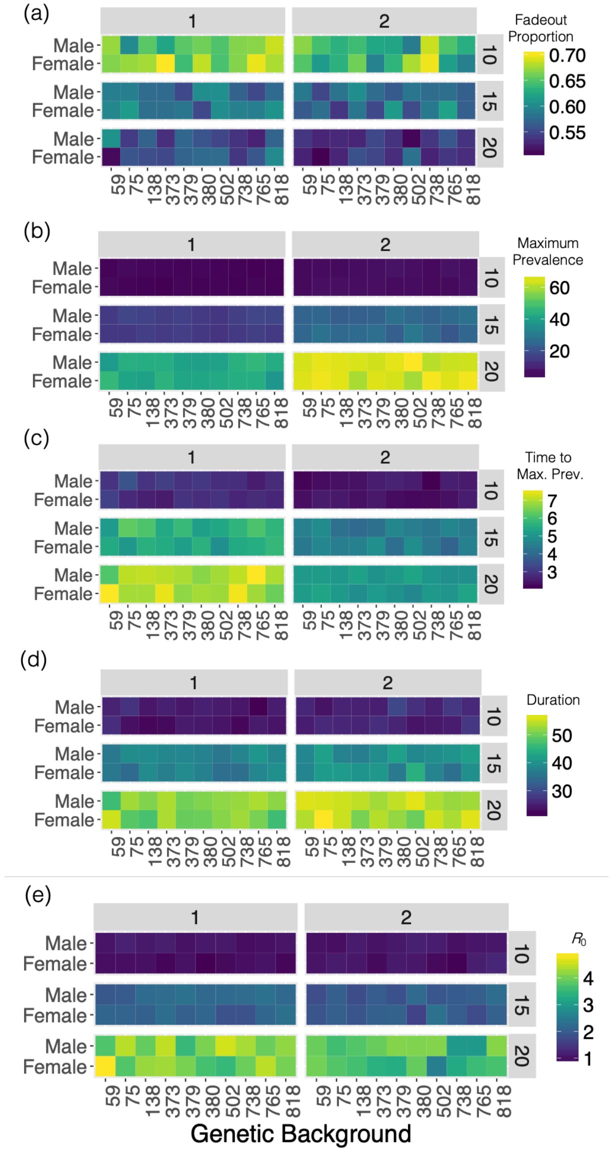


**Figure S6. Theoretical experiment #2 (τ=1.0)** - Mean (a) fadeout proportion, (b) maximum number of infected individuals (maximum prevalence), (c) the number of time steps taken to reach the maximum number of infected individuals (time to Max. Prev.) (d) the number of time steps outbreaks lasted for (duration) and (e) the number of secondary infections caused by the first infected individual ($R_0$) of outbreaks simulated in theoretical populations in experiment #2. Theoretical populations were comprised of 50 individuals from each combination of sex and genetic background. Across these simulations, pathogen transmission efficiency (τ) was 1.0, infectiousness (η) was scaled by 1 or 2 (x-axis facets) and contact network connections were based on flies aggregating within 10, 15 or 20mm of one another (r; y-axis facets).

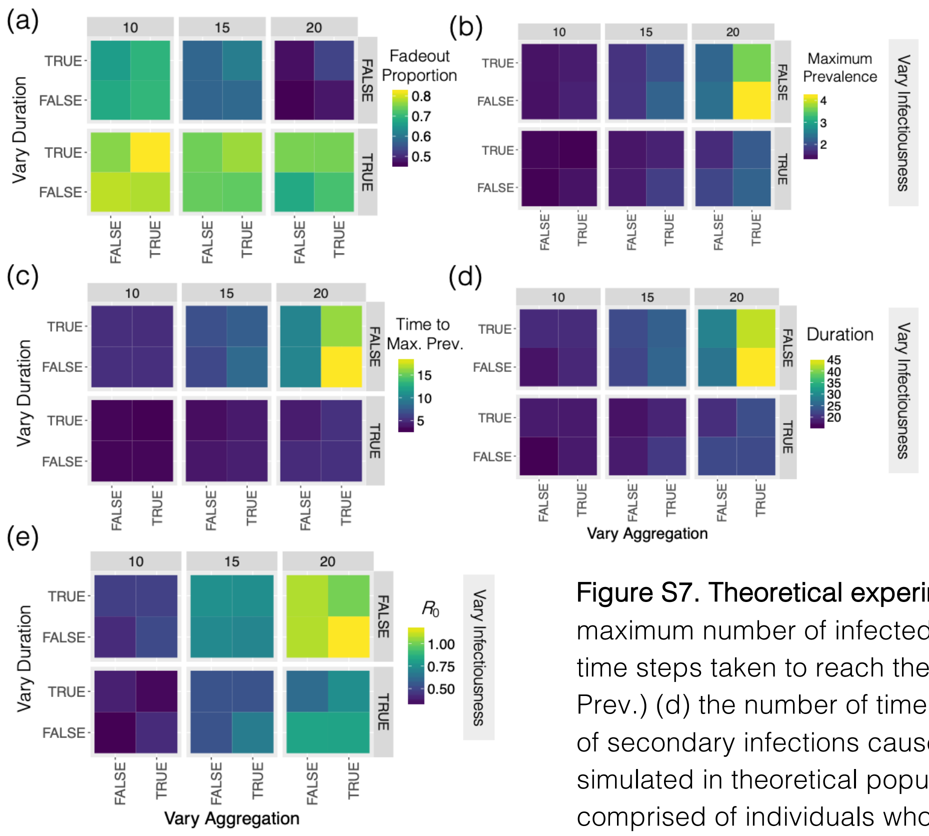


**Figure S7. Theoretical experiment #3 (τ=0.1, η=1)** - Mean (a) fadeout proportion, (b) maximum number of infected individuals (maximum prevalence), (c) the number of time steps taken to reach the maximum number of infected individuals (time to Max. Prev.) (d) the number of time steps outbreaks lasted for (duration) and (e) the number of secondary infections caused by the first infected individual ($R_0$) of outbreaks simulated in theoretical populations in experiment #3. Theoretical populations were comprised of individuals whose virus shedding, social aggregation and lifespan were derived from the variation seen across all genetic backgrounds and both sexes. The extent of this variation was manipulated by constraining variation in social aggregation (vary aggregation; x-axis), lifespan following infection (vary duration; y-axis) and virus shedding (vary infectiousness; y-axis facets) to the population mean. Across these simulations, pathogen transmission efficiency (τ) was 0.1, infectiousness (η) was 1, and contact network connections were based on flies aggregating within 10, 15 or 20mm of one another (r; x-axis facets).

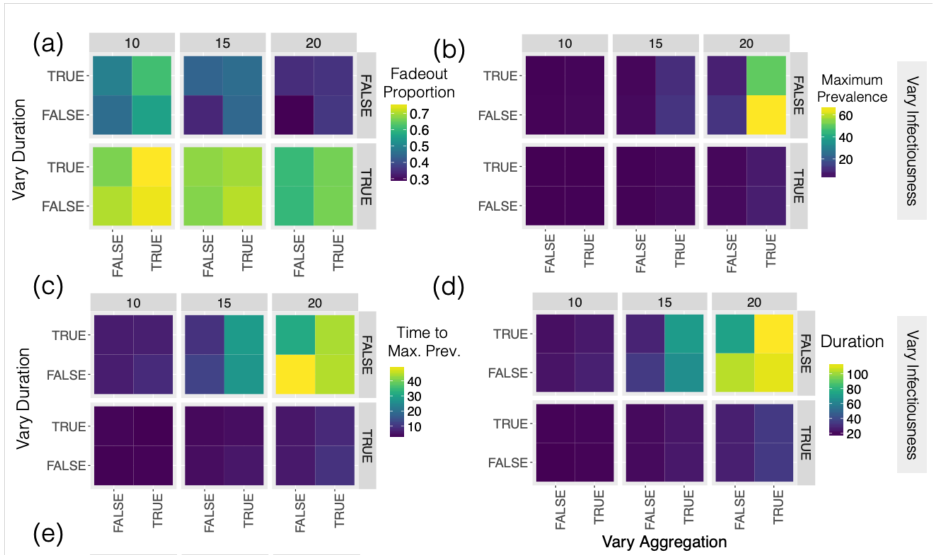


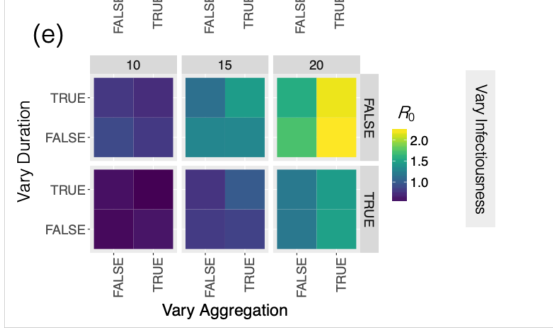


**Figure S8. Theoretical experiment #3 (τ=0.1, η=2)** - Mean (a) fadeout proportion, (b) maximum number of infected individuals (maximum prevalence), (c) the number of time steps taken to reach the maximum number of infected individuals (time to Max. Prev.) (d) the number of time steps outbreaks lasted for (duration) and (e) the number of secondary infections caused by the first infected individual ($R_0$) of outbreaks simulated in theoretical populations in experiment #3. Theoretical populations were comprised of individuals whose virus shedding, social aggregation and lifespan were derived from the variation seen across all genetic backgrounds and both sexes. The extent of this variation was manipulated by constraining variation in social aggregation (vary aggregation; x-axis), lifespan following infection (vary duration; y-axis) and virus shedding (vary infectiousness; y-axis facets) to the population mean. Across these simulations, pathogen transmission efficiency (τ) was 0.1, infectiousness (η) was 2, and contact network connections were based on flies aggregating within 10, 15 or 20mm of one another (r; x-axis facets).

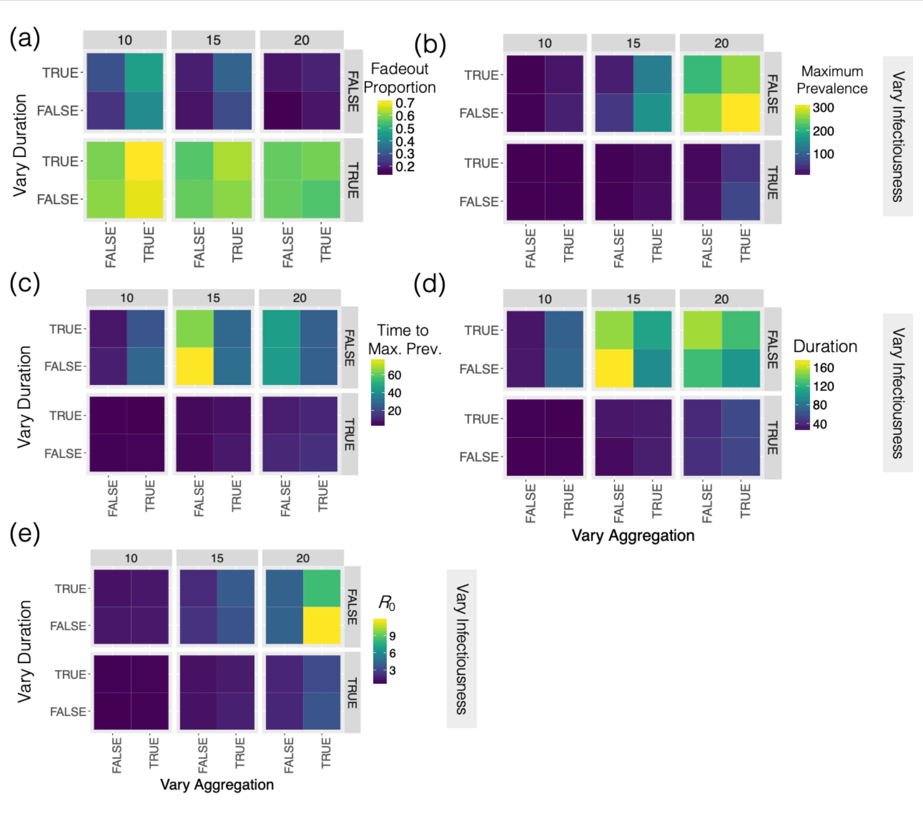


**Figure S9. Theoretical experiment #3 (τ=0.5, η=1)** - Mean (a) fadeout proportion, (b) maximum number of infected individuals (maximum prevalence), (c) the number of time steps taken to reach the maximum number of infected individuals (time to Max. Prev.) (d) the number of time steps outbreaks lasted for (duration) and (e) the number of secondary infections caused by the first infected individual ($R_0$) of outbreaks simulated in theoretical populations in experiment #3. Theoretical populations were comprised of individuals whose virus shedding, social aggregation and lifespan were derived from the variation seen across all genetic backgrounds and both sexes. The extent of this variation was manipulated by constraining variation in social aggregation (vary aggregation; x-axis), lifespan following infection (vary duration; y-axis) and virus shedding (vary infectiousness; y-axis facets) to the population mean. Across these simulations, pathogen transmission efficiency (τ) was 0.5, infectiousness (η) was 1 and contact network connections were based on flies aggregating within 10, 15 or 20mm of one another (r; x-axis facets).

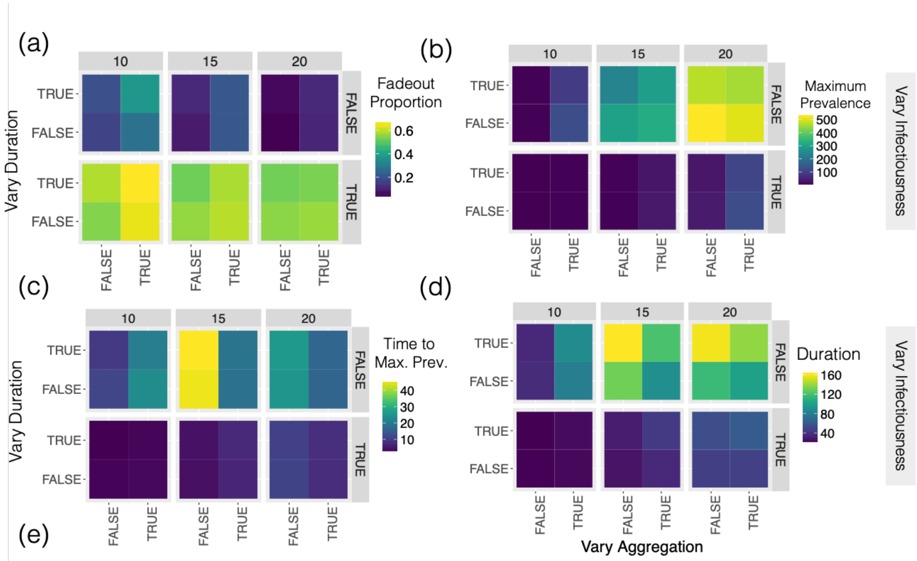


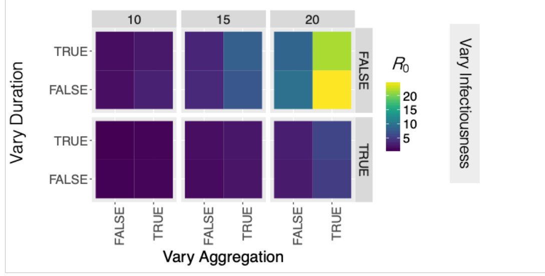


**Figure S10. Theoretical experiment #3 (τ=0.5, η=2)** - Mean (a) fadeout proportion, (b) maximum number of infected individuals (maximum prevalence), (c) the number of time steps taken to reach the maximum number of infected individuals (time to Max. Prev.) (d) the number of time steps outbreaks lasted for (duration) and (e) the number of secondary infections caused by the first infected individual ($R_0$) of outbreaks simulated in theoretical populations in experiment #3. Theoretical populations were comprised of individuals whose virus shedding, social aggregation and lifespan were derived from the variation seen across all genetic backgrounds and both sexes. The extent of this variation was manipulated by constraining variation in social aggregation (vary aggregation; x-axis), lifespan following infection (vary duration; y-axis) and virus shedding (vary infectiousness; y-axis facets) to the population mean. Across these simulations, pathogen transmission efficiency (τ) was 0.5, infectiousness (η) was 2 and contact network connections were based on flies aggregating within 10, 15 or 20mm of one another (r; x-axis facets).

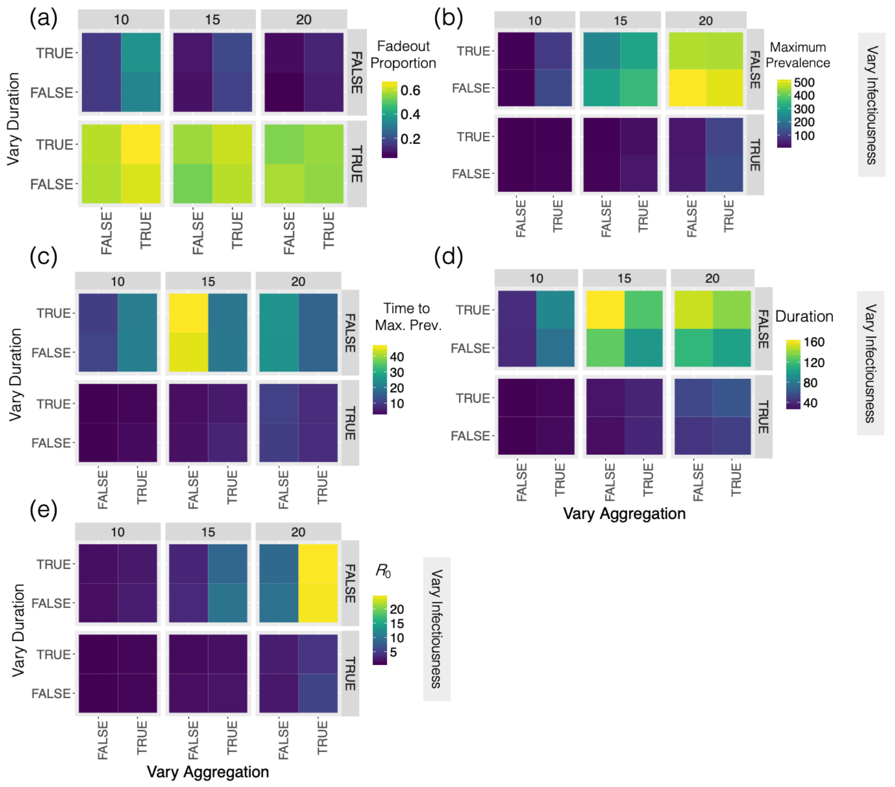


**Figure S11. Theoretical experiment #3 (τ=1.0, η=1)** - Mean (a) fadeout proportion, (b) maximum number of infected individuals (maximum prevalence), (c) the number of time steps taken to reach the maximum number of infected individuals (time to Max. Prev.) (d) the number of time steps outbreaks lasted for (duration) and (e) the number of secondary infections caused by the first infected individual ($R_0$) of outbreaks simulated in theoretical populations in experiment #3. Theoretical populations were comprised of individuals whose virus shedding, social aggregation and lifespan were derived from the variation seen across all genetic backgrounds and both sexes. The extent of this variation was manipulated by constraining variation in social aggregation (vary aggregation; x-axis), lifespan following infection (vary duration; y-axis) and virus shedding (vary infectiousness; y-axis facets) to the population mean. Across these simulations, pathogen transmission efficiency (τ) was 1.0, infectiousness (η) was 1 and contact network connections were based on flies aggregating within 10, 15 or 20mm of one another (r; x-axis facets).

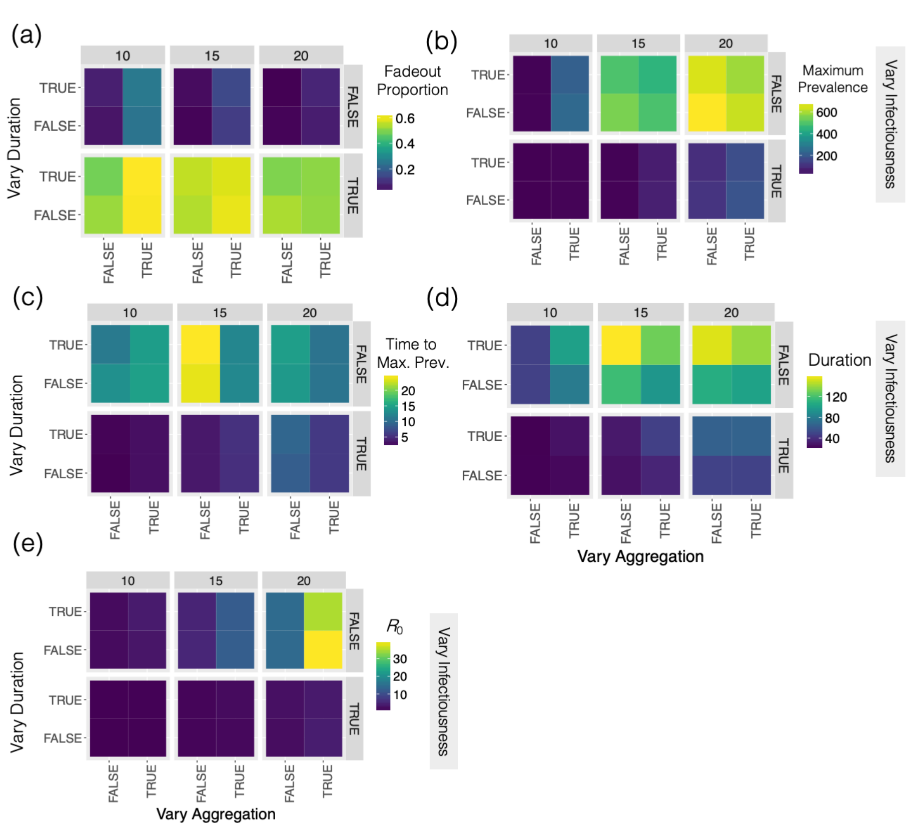


**Figure S12. Theoretical experiment #3 (τ=1.0, η=2)** - Mean (a) fadeout proportion, (b) maximum number of infected individuals (maximum prevalence), (c) the number of time steps taken to reach the maximum number of infected individuals (time to Max. Prev.) (d) the number of time steps outbreaks lasted for (duration) and (e) the number of secondary infections caused by the first infected individual ($R_0$) of outbreaks simulated in theoretical populations in experiment #3. Theoretical populations were comprised of individuals whose virus shedding, social aggregation and lifespan were derived from the variation seen across all genetic backgrounds and both sexes. The extent of this variation was manipulated by constraining variation in social aggregation (vary aggregation; x-axis), lifespan following infection (vary duration; y-axis) and virus shedding (vary infectiousness; y-axis facets) to the population mean. Across these simulations, pathogen transmission efficiency (τ) was 1.0, infectiousness (η) was 2 and contact network connections were based on flies aggregating within 10, 15 or 20mm of one another (r; x-axis facets).

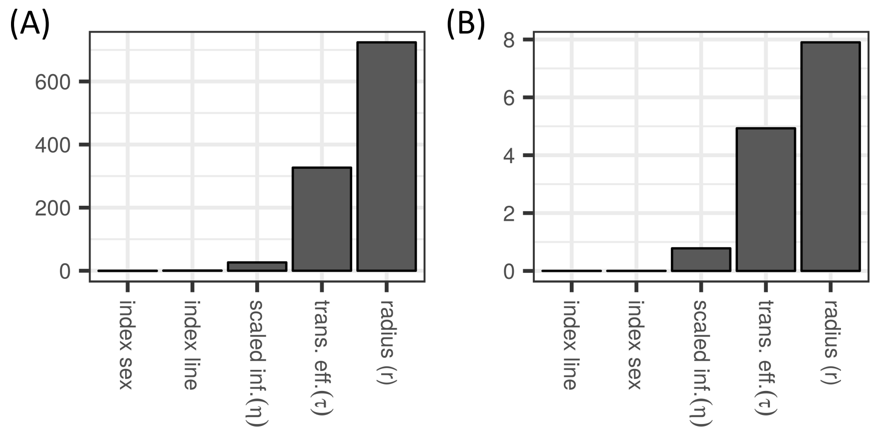


**Figure S13. Theoretical Experiment #2, (r=15, τ=1.0, η=2).** Results of variable importance analysis for Experiment #2 with n=1000 trees using *cforest* function from the *party* package in R. Variables are listed on x-axis. Y-axis describes variable importance (mean decrease in accuracy [MDA]). Which variables most determine (A) how long an infection lasts, and (B) the number of secondary cases of infection caused by the first individual to become infected?
